# Multifaceted roles of H2B mono-ubiquitylation in D-loop metabolism during homologous recombination repair

**DOI:** 10.1101/2024.09.13.612919

**Authors:** Shih-Hsun Hung, Yuan Liang, Wolf-Dietrich Heyer

## Abstract

Repairing DNA double-strand breaks is crucial for maintaining genome integrity, which occurs primarily through homologous recombination (HR) in *S. cerevisiae.* Nucleosomes, composed of DNA wrapped around a histone octamer, present a natural barrier to end-resection to initiate HR, but the impact on the downstream HR steps of homology search, DNA strand invasion and repair synthesis remain to be determined. Displacement loops (D-loops) play a pivotal role in HR, yet the influence of chromatin dynamics on D-loop metabolism remains unclear. Using the physical D-loop capture (DLC) and D-loop extension (DLE) assays to track HR intermediates, we employed genetic analysis to reveal that H2B mono-ubiquitylation (H2Bubi) affects multiple steps during HR repair. We infer that H2Bubi modulates chromatin structure, not only promoting histone degradation for nascent D-loop formation but also stabilizing extended D-loops through nucleosome assembly. Furthermore, H2Bubi regulates DNA resection *via* Rad9 recruitment to suppress a feedback control mechanism that dampens D-loop formation and extension at hyper-resected ends. Through physical and genetic assays to determine repair outcomes, we demonstrate that H2Bubi plays a crucial role in preventing break-induced replication and thus promoting genomic stability.

**Graphical Abstract:** 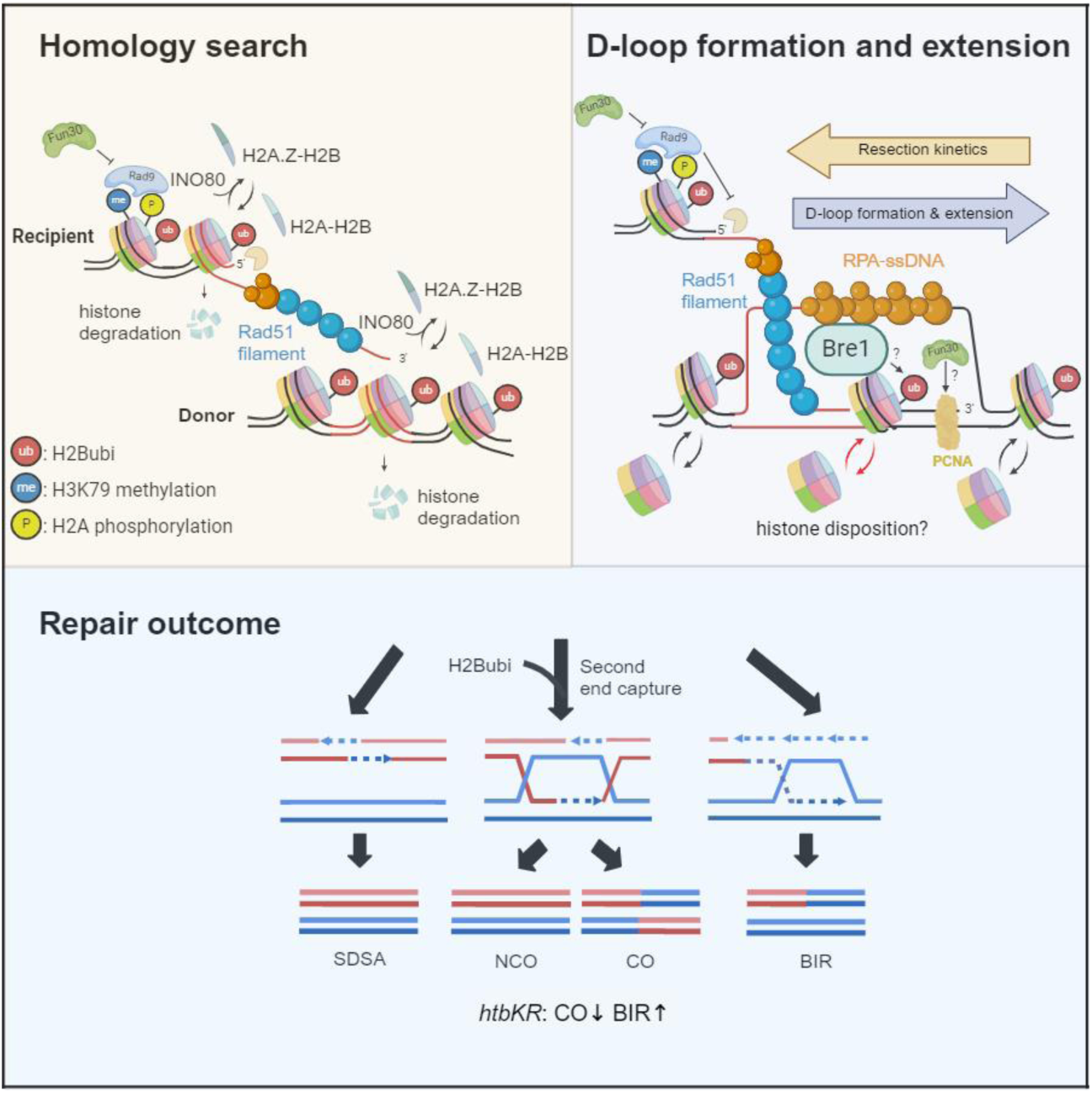

**Highlights:**

- H2Bubi is epistatic to H2A.Z and INO80 in promoting homology search and D-loop formation
- H2Bubi stabilizes extended D-loop
- Excessive resection counteracts D-loop formation and extension
- H2Bubi promotes crossover events and limits the frequency of break-induced replication outcomes in HR repair

## INTRODUCTION

Homologous recombination (HR) repair stands as a primary mechanism for resolving DNA double-strand breaks (DSBs) in *Saccharomyces cerevisiae*. During this process, the break termini undergo resection, leading to the formation of 3’ single-strand DNA (ssDNA) tails, a crucial step in committing the DNA break to repair *via* HR (Cejka and Symington, 2021; Symington, 2014). The recombinase protein Rad51 assembles onto the ssDNA to form the Rad51-ssDNA presynaptic filament, which is capable of searching for homology and invading homologous DNA, ultimately forming a DNA joint molecule termed the displacement loop (D-loop) (Heyer et al., 2010). DNA synthesis subsequently commences from the 3’ end of the invading strand, and the extended D-loop undergoes resolution through one of several sub-pathways to restore the integrity of the damaged chromosome, including synthesis-dependent strand annealing (SDSA), double-strand break repair involving double Holliday junction formation (DSBR), and break-induced replication (BIR) (Wright et al., 2018).

Two steps of HR undergo reversal and exist in a dynamic balance between forward and backward reactions (Heyer, 2015; Piazza and Heyer, 2019): the Rad51–ssDNA filament and the nascent D-loop. The formation of the Rad51–ssDNA filament requires the assistance of mediator proteins, while the Srs2 helicase/translocase disrupts the Rad51–ssDNA filament (Antony et al., 2009; Burgess et al., 2009; Dupaigne et al., 2008; Krejci et al., 2004; Krejci et al., 2003; Liu et al., 2011b; Ma et al., 2018; Roy et al., 2021; Van Komen et al., 2003; Veaute et al., 2003). Additionally, the Sgs1-Top3-Rmi1 (STR) complex is involved in the disruption of the nascent D-loop (Cejka et al., 2012; Fasching et al., 2015; Lo et al., 2006; Piazza et al., 2019; Wu and Hickson, 2003). Both Srs2 and Mph1 participate in the disruption of extended D-loops (Liu et al., 2017; Mitchel et al., 2013; Piazza et al., 2019; Prakash et al., 2009). Chromatin structure has been conceptualized as a barrier to repair that requires lifting and subsequent restoration (Polo and Almouzni, 2015). Chromatin regulators have been shown to affect HR repair, especially end resection (Chakraborty et al., 2021; Garcia Fernandez and Fabre, 2022; Hauer and Gasser, 2017; Mohan et al., 2021; Papamichos-Chronakis and Peterson, 2013; Peterson and Almouzni, 2013; Price and D’Andrea, 2013; Seeber et al., 2013b; Seeber et al., 2018; Smeenk and van Attikum, 2013; van Attikum and Gasser, 2009). However, understanding of how chromatin regulators influence the kinetics of D-loop metabolism has been limited due to the absence of assays capable detecting nascent and extended D-loops during DNA repair in mitotically growing cells. Using a proximity ligation approach, this gap in the toolbox to study HR-intermediates has been closed by the development of the D-loop capture (DLC) and D-loop extension (DLE) assays (Piazza et al., 2018; Piazza et al., 2019). This progress allowed to define two pathways of D-loop disruption, one involving the STR complex, likely favoring nascent D-loops, and one involving Srs2 involving extended, possibly longer D-loops (Piazza et al., 2019). Interestingly, the STR complex acts epistatically to Mph1, which is also capable of disrupting extended D-loops (Piazza et al., 2019; Prakash et al., 2009). Unlike other PCR-based approaches to study D-loop extension, which measure the dsDNA repair product (Donnianni and Symington, 2013; Jain et al., 2009), the DLE-assay allows to track the extension intermediate before conversion to dsDNA product (Piazza et al., 2018). Collectively, these advances provide avenues for studying D-loop metabolism within the context of chromatin.

In eukaryotes, DNA intricately wraps around histones to form nucleosomes. Thus, dynamic alterations in chromatin structure during HR repair can profoundly affect DNA resection, the subsequent homology search, followed by D-loop formation, and extension. Chromatin structure is regulated through several mechanisms: histone modification, non-histone proteins, histone variants, histone chaperones, and ATP-dependent chromatin-remodeling enzymes (Papamichos-Chronakis and Peterson, 2013). In eukaryotic DNA replication, nucleosome disruption ahead of the replication fork and subsequent reassembly behind are essential processes (Groth et al., 2007a). During D-loop extension, which resembles lagging strand DNA replication, the involvement of the DNA replication machinery including DNA polymerase delta, PCNA and RFC is necessary (Donnianni and Symington, 2013; Li et al., 2009; Liu et al., 2021b; Lydeard et al., 2007; McVey et al., 2016; Reitz et al., 2023; Sneeden et al., 2013). However, whether nucleosomes are assembled into the three-strand stage of the D-loop remains an open question. Histone chaperones such as Asf1 and CAF-1 play crucial roles in regulating chromatin replication (Groth et al., 2007b; Ransom et al., 2010). They have also been reported to stabilize D-loops at stalled replication forks, ensuring replication restart in fission yeast (Hardy et al., 2019; Pietrobon et al., 2014). Moreover, nucleosome assembly at D-loops was inferred from studies of the HR-effect of human ATRX (Juhasz et al., 2018).

Mono-ubiquitylation of H2B at lysine 123 (H2Bubi) has a profound impact on chromatin structure interfering with chromatin compaction resulting in an open and accessible conformation (Fierz et al., 2011). Rad6, in cooperation with the E3 enzyme Bre1, plays a pivotal role in regulating H2Bubi in *Saccharomyces cerevisiae*. H2Bubi has been associated with DSB repair through relaxing chromatin structure, as observed in studies conducted in both *Saccharomyces cerevisiae* and human cell lines (Challa et al., 2021; Moyal et al., 2011; Nakamura et al., 2011; Xu et al., 2016; Yamashita et al., 2004; Zheng et al., 2018). H2Bubi is reported to affect H3 methylation through Dot1-mediated H3K79 and Set1-mediated H3K4 methylation (Game and Chernikova, 2009). However, previous studies suggest that the HR repair function of H2Bubi is independent of H3K4 or H3K79 methylation (Zheng et al., 2018). Moreover, H2Bubi helps avoid hyper-resection by permitting Dot1-dependent H3K79 methylation for Rad9 recruitment, which is known to limit excessive resection (Fierz et al., 2011; Giannattasio et al., 2005; Zheng et al., 2018). Despite this, the role of H2Bubi in fine-tuning chromatin structure for DSB repair presents a paradox: it enhances accessibility for repair while simultaneously maintaining Rad9 chromatin binding to restrict excessive DNA end resection, thereby limiting the availability of single-stranded DNA (ssDNA) necessary for strand invasion. Investigating how H2Bubi affects DNA resection and D-loop metabolism should elucidate the molecular mechanisms underlying the complex role of H2Bubi in HR repair.

Here, we unveil the multi-faceted roles of H2Bubi in D-loop metabolism and its effect on DSB repair pathway usage in *S. cerevisiae*. Through epistasis analysis using the DLC and DLE assays to determine D-loop levels and their extension, we reveal that H2Bubi modulates chromatin structure, not only promoting histone degradation for nascent D-loop formation but also stabilizing extended D-loops through nucleosome assembly. Furthermore, H2Bubi regulates DNA resection *via* Rad9 recruitment, suppressing a feedback control mechanism that dampens D-loop formation and extension. Through physical and genetic assays aimed at studying repair outcomes, we demonstrate that H2Bubi plays a crucial role in preventing break-induced replication, a sub-pathway of DNA double-strand break repair known to culminate in genomic instability.

## RESULTS

### H2Bubi affects the kinetics of D-loops metabolism

To directly investigate the involvement of H2Bubi in the formation and extension of D-loops during HR repair, we employed the DLC and DLE assays (Piazza et al., 2018; Piazza et al., 2019; Reitz et al., 2022), which are able to quantify the kinetics of nascent D-loop intermediate formation and extension *in vivo* and distinguish themselves from other assays limited to measuring the final physical or genetic repair end products.

A site-specific double-strand break (DSB) induced by HO endonuclease on chromosome V leads to homology search and DNA strand invasion of a 2 kb homologous donor sequence on chromosome II. Two *EcoR*1 sites were used for proximity ligation (**Figure 1B**). After *GAL-HO* induction and DNA strand invasion at the donor site, *in vivo* inter-strand DNA crosslinking by psoralen covalently links the heteroduplex DNA (hDNA) within the D-loop, preserving it during subsequent steps. A long complementary oligonucleotide is utilized to restore the restriction site that was ablated by DNA end resection. Subsequently, following restriction site restoration and digestion, the crosslinked hDNA, held together by psoralen, is preferentially ligated during the proximity ligation reaction under dilute conditions. The resulting unique chimeric ligation products are quantified by quantitative PCR (qPCR) using a pair of specific primers, and this PCR product is referred to as the DLC signal (**Figure 1A**). Extended D-loops were quantified using a similar proximity ligation approach to examine D-loop extension to the downstream 396 base-pair *Hind*III site in the absence of psoralen crosslinking (**Figure 1E**). Two long complementary oligonucleotides are used to restore the restriction sites. One serves to restore the restriction site ablated by DNA end resection, while the other is used to restore the *Hind*III site located 396 base-pair downstream of the invading 3’-OH end in the ssDNA extension intermediate in unique sequence following the homologous donor sequence. Following restriction site restoration and digestion, proximity ligation is employed to ligate the extended invading strand into a stable circular form of DNA molecule. The resulting unique chimera is quantified by qPCR using specific primers, and this PCR product is referred to as the DLE signal (**Figure 1D**). The DLC and DLE assays contain a number of quantitative controls to ensure reproducible quantitation including monitoring the level of DSB formation, a normalization control (*ARG4* locus), as well as controlling for the intramolecular ligation efficiency by measuring the ligation of a circularized DNA fragment from chromosome VIII following *Eco*R1 digestion (DLC assay) and a circularized DNA fragment from chromosome XII following *Hind*III digestion (DLE assay) (Piazza et al., 2018; Piazza et al., 2019; Reitz et al., 2022).

**Figure 1.**
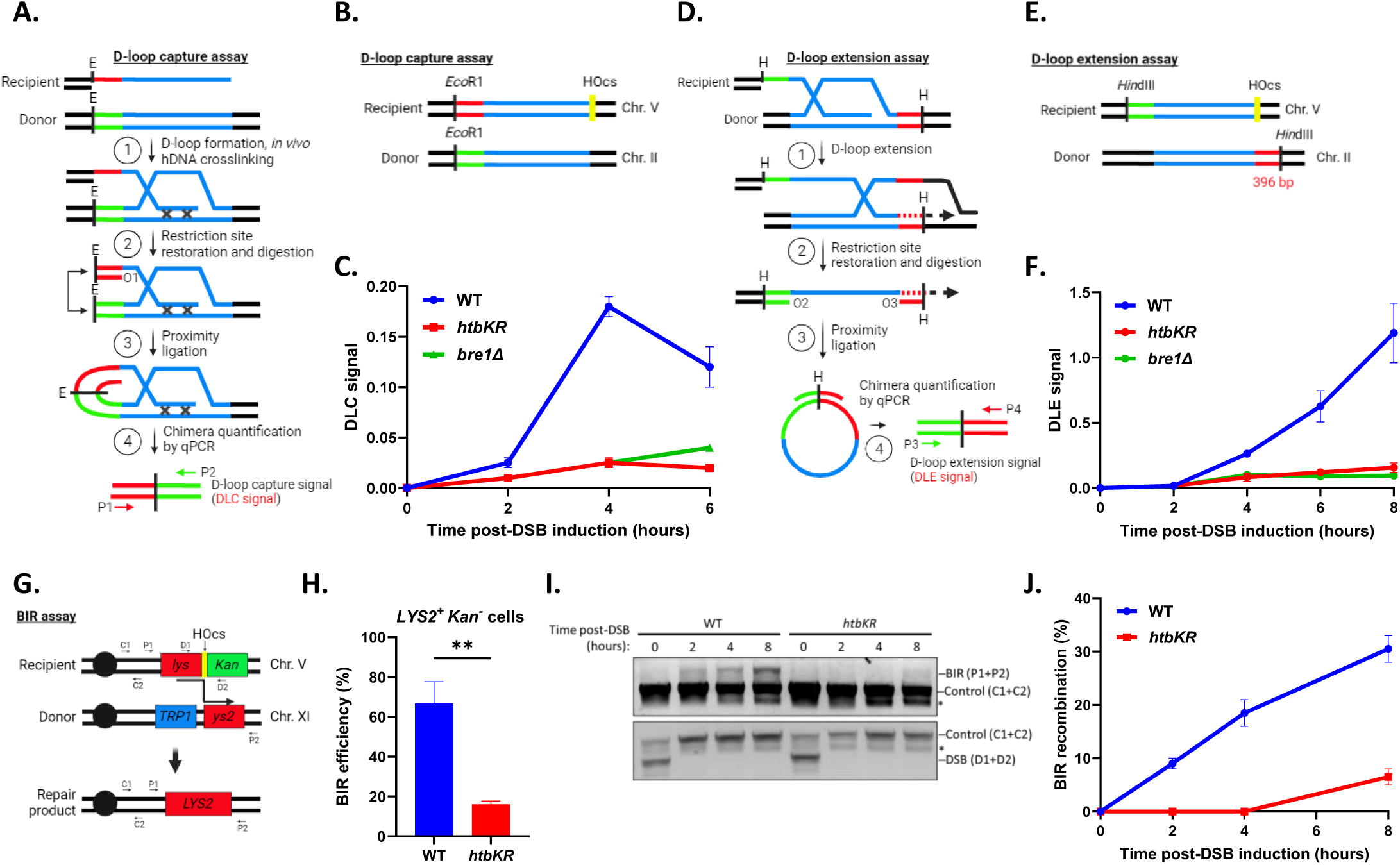
H2Bubi affects D-loop metabolism. (**A**) Schematic representation illustrating the rationale behind the D-loop capture (DLC) assay (See text for details). (**B**) Schematic representation depicting the construct of the DLC assay strain. (**C**) Quantification of the DLC signal from the specified strains obtained at various time points following HO induction. Error bars, SEM (n = 2). (**D**) Schematic representation illustrating the rationale behind the D-loop extension (DLE) assay (Refer to the text for details). (**E**) Schematic representation depicting the construct of the DLE assay strain. (**F**) Analysis of the DLE signal from the specified strains was conducted at various time points following HO induction. Error bars, SEM (n ≥ 2). (**G**) Schematic representation illustrating the construct for the Break-Induced Replication (BIR) assay (See text for details) (**H**) BIR efficiency was determined by counting colony formation on YP-Gal (ensure that colonies are viable on LYS drop-out (*LYS2+*) and sensitive to geneticin (*Kan-*)) divided by YP-Glu for each of the indicated strains across three independent trials. Error bars, SEM (n = 3), **: P=0.0014. (**I**) Agarose gel analysis was conducted to assess PCR products from various primer sets, examining the kinetics of Break-Induced Replication (BIR) product formation (P1 and P2), DNA loading control (C1 and C2), and DSB cleavage efficiency (D1 and D2) from the indicated strains at various time points following HO induction. (**J**) Plot illustrating the quantification of Break-Induced Replication (BIR) products at each time point, as obtained from (**I**).

As anticipated from previous results (Piazza et al., 2018; Piazza et al., 2019; Reitz et al., 2022), in wild-type (WT) cells, D-loop formation was observed around 2 hours post DSB induction, peaking at 4 hours before slightly declining at 6 hours due to D-loop extension and migration (**Figure 1C**). As expected, D-loop extension exhibited slower kinetics, first detected after 4 hours, and gradually increasing until 8 hours (**Figure 1F**). Nascent and extended D-loops could be investigated separately due to this 2-hour difference (Piazza et al., 2018; Piazza et al., 2019). We observed a significant decrease in DLC signal and DLE signal at all time points upon deletion of *BRE1* (*bre1*Δ) or in the *htbKR* mutant (lacking monoubiquitylation of H2B on lysine (K) 123 due to a amino acid change to arginine (R)) (**Figures 1C** and **1F**). The DLC signal at 4 hours post-DSB in the *bre1*Δ and *htbKR* mutants resembled the DLC signal at 2 hours post-DSB in WT cells but with no significant increase thereafter. Additionally, the DLE signal was significantly depressed with essentially no increase after 4 hours post DSB-induction in the *bre1*Δ and *htbKR* mutant cells. Control experiments showed similar level of DSB induction efficiency by HO endonuclease and consistent individual values for the DLC and DLE control experiments in WT, *bre1*Δ and *htbKR* cells (**Figure S1**). These results suggest that H2Bubi promotes DNA strand invasion to form and/or maintain nascent and extended D-loops. While the defect in D-loop extension could be a mere consequence of the defect in D-loop levels, additional genetic analysis presented below suggests that both defects reflect separate functions of H2Bubi in HR employing distinct mechanisms.

The DLE assay is designed to detect the extension intermediate of BIR, as the system does not provide homology on the other DSB end to make the DSB repairable. This design was implemented to avoid complications from cells that repaired the DSB and resumed growth (Piazza et al., 2018; Piazza et al., 2019; Reitz et al., 2022). To further investigate the impact of H2Bubi on the kinetics of repair product formation, we used a well-characterized genetic and physical assay for BIR (Donnianni and Symington, 2013; Donnianni et al., 2019). The repair of an induced HO cut on chromosome V depends on the search for and invasion of homology on chromosome XI (**Figure 1G**). Following DSB induction, successful homology-directed repair can only occur *via* BIR to restore a functional *LYS2* gene, as evidenced by colony growth on synthetic medium lacking lysine. The loss of the *KanMX* gene confirms that the DSB was repaired by BIR and not by non-homologous end joining (**Figure 1H**). BIR efficiency was assessed by measuring colony formation after DSB induction (**Figure 1H**), and BIR kinetics were evaluated by detecting the BIR repair intermediates/products *via* PCR (**Figures 1I** and **1J**).

We observed a significant decrease in BIR efficiency in the *htbKR* mutant (**Figure 1H**). Furthermore, BIR kinetics in the *htbKR* mutant was markedly delayed, emerging only at 8 hours post-DSB induction, compared to the WT cells, where BIR products appeared 2 hours post-DSB induction and gradually increased over time (**Figures 1I and J**). The disparity in BIR kinetics between WT and *htbKR* mutants does not seem to stem from differences in HO induction efficiency, as evidenced by comparable cutting efficiencies at the HO cut site. Significantly, no band indicative of an intact HO cut site was detected 2 hours post-HO induction in both WT and *htbKR* mutant, as revealed by PCR, indicating near 100% DSB induction (**Figure 1I**). Collectively, these findings strongly suggest that H2Bubi affects the kinetics of D-loop metabolism.

### D-loop disruption pathways counteract nascent D-loop formation in the absence of H2Bubi

To further elucidate the role of H2Bubi in the dynamic equilibrium of nascent D-loop formation and disruption, we aimed to measure the DLC signal at 2 hours post-DSB induction. At this time point, D-loop extension is very limited, allowing for the specific capture of nascent D-loops (Piazza et al., 2018; Piazza et al., 2019). It has been reported that two distinct pathways, one STR- and Mph1-dependent and the other Srs2-dependent, target different D-loop species (**Figure 2A**) (Piazza et al., 2019). We thus compared the DLC signal in *htbKR* mutant in combination with *SGS1* (*sgs1*Δ) or *SRS2* (*srs2*Δ) deletions. In line with our previous results (Piazza et al., 2019), we observed that *sgs1*Δ or *srs2*Δ resulted in a significant 2- to 3-fold increase in DLC signal (**Figures 2B** and **2C**). In addition, we observed that the reduced level of DLC signal in the *htbKR* mutant was restored to WT levels when combined with the *SGS1* deletion, although it remained lower than in the *sgs1*Δ single mutant (**Figure 2B**). Surprisingly, in the *srs2*Δ *htbKR* double mutant, the DLC signal was restored to a level similar to that of the *srs2*Δ single mutant (**Figure 2C**). To delve deeper into the epistatic relationship between H2Bubi and *SRS2*, we disabled the function of the STR complex by overexpressing a catalytically inactive mutant *top3-Y356F* (*top3-cd*) in both the *srs2*Δ single mutant and *srs2*Δ *htbKR* double mutant backgrounds along with wild type Top3 as a control. This approach to generate a double mutant context is necessary, as the *sgs1*Δ *srs2*Δ double mutant is synthetically lethal (Fabre et al., 2002). Our findings revealed a significant 2-fold increase in the DLC signal when *top3-cd* was overexpressed in the *srs2*Δ mutant background, consistent with the previous conclusion that the Srs2- and STR-dependent pathways are separate (Piazza et al., 2019). However, no significant change was observed upon overexpression of either Top3 or *top3-cd* in the *srs2*Δ *htbKR* double mutant background (**Figure 2D**). Together, these results suggest that H2Bubi promotes nascent D-loop levels. Notably, the deficiency in nascent D-loop levels can be restored by eliminating STR- or Srs2-dependent D-loop disruption pathways (Piazza et al., 2019), underscoring the critical role of the D-loop disruption pathways in monitoring nascent D-loop levels. Moreover, H2Bubi may preferentially promote longer D-loop formation, which was proposed to be the target of the Srs2-dependent pathway (**Figure 2E**). Alternatively, Rad51-ssDNA filament stabilized by *srs2*Δ may promote homology search thus compensate the nascent D-loop formation defect in the *htbKR* mutant. We consider this alternative less likely, as the Rad51 paralog complex Rad55-Rad57 effectively insulates Rad51-sDNA filaments against disruption by Srs2 during DSB repair (Liu et al., 2011a).

**Figure 2.**
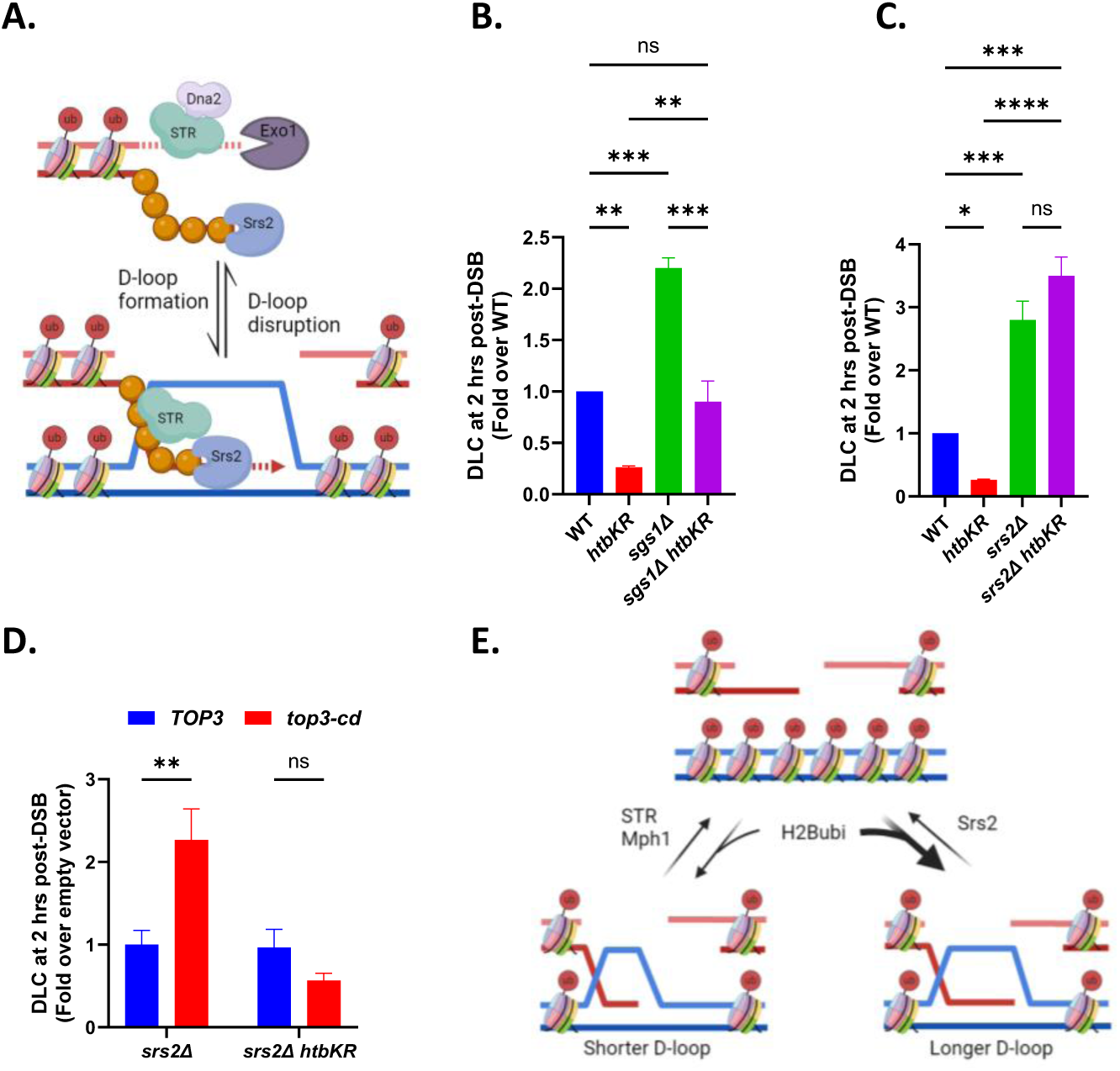
D-loop disruption pathways prohibit D-loop formation in the absence of H2Bubi. (**A**) Homologous recombination repair begins with the invasion of a homologous duplex by a 3’ overhang formed through DNA end resection (Sgs1-Top3-Rmi1 (STR)-Dna2 or Exo1). The invasion of the 3’ end is monitored by the STR and Mph1 epistatic pathways, alongside the Srs2-dependent D-loop disruption pathway. (**B**) Quantification of the DLC signal in WT, *htbKR*, and *sgs1Δ* was examined individually and in double mutant combinations. Error bars, SEM (n ≥ 2). **: P= 0.0068, ***: P= 0.0002. (**C**) Quantification of the DLC signal in WT, *htbKR* and *srs2*Δ was examined individually or in double mutant combinations. Error bars, SEM (n ≥ 2). *: P= 0.04, ***: P= 0.0008, ****: P= 0.0001, ns: not significant. (**D**) Quantification of the DLC signal in the *htbKR* mutant, either with the absence of Srs2 or with the absence of both STR and Srs2-dependent D-loop disruption pathways. Error bars, SEM (n = 3). **: P=0.006, ns: not significant. (**E**) H2Bubi may promote longer D-loop formation, as evidenced by the observation that blocking the Srs2-dependent pathway rescues more D-loop levels compared to blocking the STR pathway.

### H2Bubi is epistatic to H2A.Z and INO80 in promoting nascent D-loop formation

Checkpoint activation and the INO80 chromatin remodeling complex (INO80-C) trigger histone degradation through the recruitment of several ubiquitin ligases and the proteasome to damaged chromatin, resulting in reduced chromatin compaction and enhanced chromosome movement (Challa et al., 2021; Cheblal et al., 2020; Hauer et al., 2017; Seeber et al., 2013a). We thus determined the DLC signal in mutants in two well-studied chromatin regulators, the INO80 complex (INO80-C) and the high-mobility group proteins *NHP6A/B*, which were previously described as having compacted and expanded chromatin, respectively (Hauer et al., 2017) (**Figure 3A**). We measured the DLC signal at 2 hrs, which represents nascent D-loops, as extension starts only later (**Figure 1C** and **1F**). Indeed, the deletion of the *INO80-C* subunit *ARP8* (*arp8*Δ) resulted in a 5-fold reduction of the DLC signal compared to the WT, and no significant difference was observed between *arp8*Δ, *htbKR*, and *arp8*Δ *htbKR* mutants (**Figure 3B**). In contrast, the reduced DLC signal in the *htbKR* mutant was restored to WT levels when combined with the *NHP6A/B* deletion (*nhp6*ΔΔ) mutant (**Figure 3C**). This restoration is likely due to a 20% reduction in histone occupancy in the *nhp6*ΔΔ background (Celona et al., 2011; Challa et al., 2021; Cheblal et al., 2020), indicating a role for H2Bubi in promoting nascent D-loop levels *via* histone degradation. Our results are consistent with a previous study demonstrating that Rad6 and Bre1 are both recruited to chromatin upon DNA damage and contribute to recombination by monitoring chromosome movement and the total level of repair products (Challa et al., 2021).

**Figure 3.**
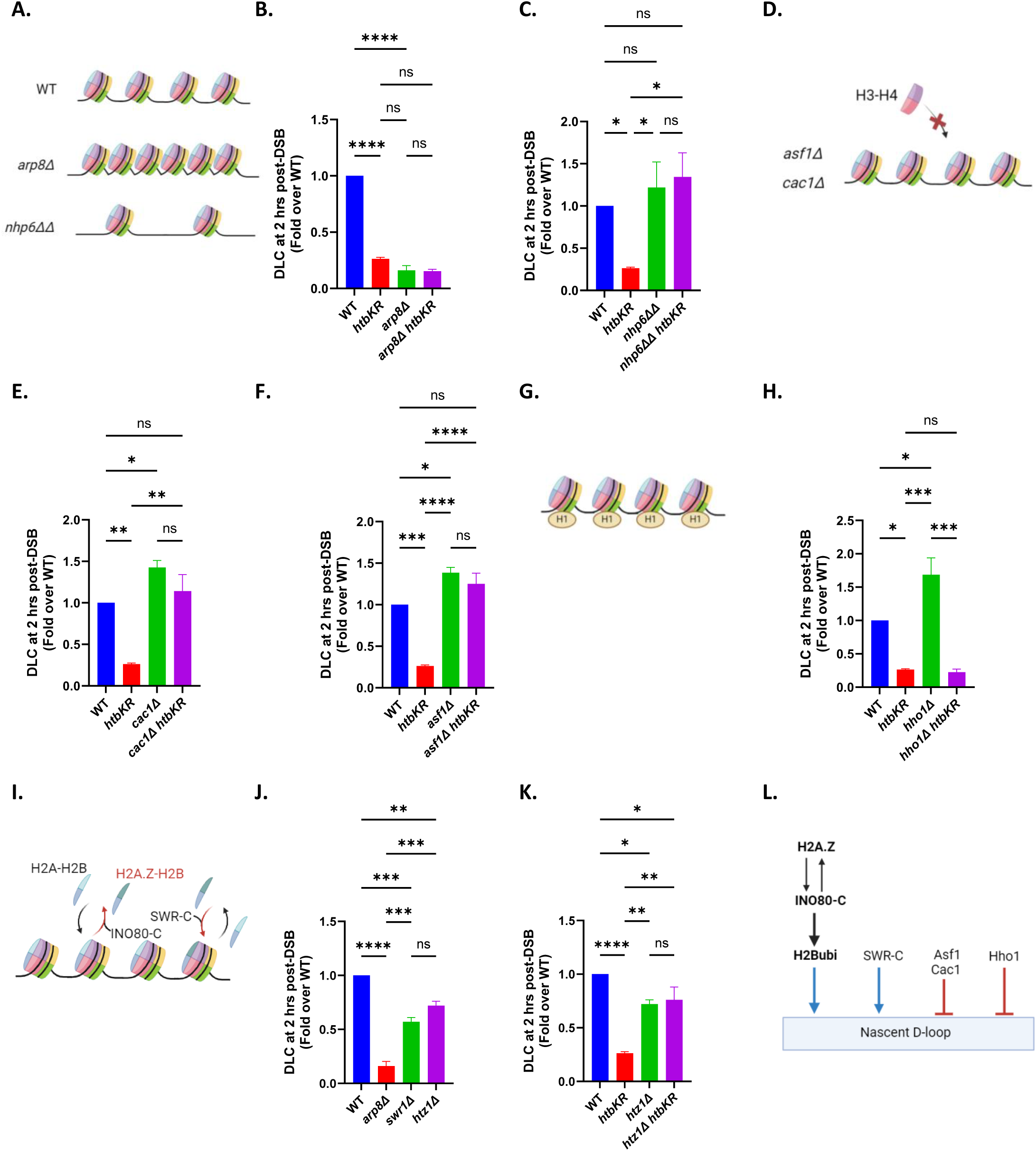
H2Bubi is epistatic to H2A.Z and INO80 in promoting nascent D-loop formation. (**A**) Schematic representation showing that *nhp6ΔΔ* cells exhibit reduced histone occupancy, while *arp8Δ* cells display more condensed chromatin. (**B**) DLC signal in WT, *htbKR*, *arp8Δ*, and *arp8Δ htbKR* strains. Error bars, SEM (n = 3). ****: P< 0.0001, ns: not significant. (**C**) DLC signal in WT, *htbKR*, *nhp6ΔΔ* and *nhp6ΔΔ htbKR* strains. Error bars, SEM (n ≥ 2). *: P< 0.03, ns: not significant. (**D**) Asf1 and CAF-1 are histone chaperones responsible for depositing histone H3-H4 onto DNA. Lack of Asf1 or CAF-1 results in reduced histone occupancy. (**E**) DLC signal in WT, *htbKR*, *cac1Δ and cac1Δ htbKR* strains. Error bars, SEM (n ≥ 2). *: P= 0.04, **: P< 0.0015, ns: not significant. (**F**) DLC signal in WT, *htbKR*, *asf1Δ and asf1Δ htbKR* strains. Error bars, SEM (n ≥ 2). *: P= 0.01, ***: P= 0.0002, ****: P< 0.0001, ns: not significant. (**G**) Histone H1 is a linker histone that binds the outside of nucleosomes and modifies chromatin structure. (**H**) DLC signal in WT, *htbKR*, *hho1Δ* and *hho1Δ htbKR* strains. Error bars, SEM (n = 3). *: P< 0.02, ***: P= 0.0002, ns: not significant. (**I**) The INO80 and SWR complexes are chromatin remodelers responsible for facilitating histone dimer exchange. (**J**) DLC signal in WT, *arp8Δ*, *swr1Δ* and *htz1Δ* strains. Error bars, SEM (n = 2). **: P= 0.005, ***: P< 0.0007, ****: P< 0.0001, ns: not significant. (**K**) DLC signal in WT, *htbKR*, *htz1Δ* and *htz1Δ htbKR* strains. Error bars, SEM (n ≥ 2). *: P< 0.04, **: P< 0.002, ****: P< 0.0001, ns: not significant. (**L**) A schematic representation indicating that H2A.Z and INO80-C likely operates upstream of H2Bubi in regulating D-loop formation. SWR1-C promotes nascent D-loop formation. Hho1, Asf1, and Cac1 negatively regulate nascent D-loop formation.

To further support the notion that H2Bubi affects HR through chromatin decompaction, we examined the histone chaperons CAF-1 and Asf1, which are involved in chromatin assembly through histone H3-H4 deposition (Tyler et al., 1999) (**Figure 3D**). We used a *cac1*Δ strain to disable CAF-1 and the *asf1*Δ mutant. Both exhibited a 1.5-fold increase in nascent D-loop level compared to WT cells (**Figure 3E** and **3F**), consistent with the role of Asf1 and CAF-1 in preventing unwanted recombination through the deposition of histone H3-H4 (Prado et al., 2004). Additionally, deletions of *ASF1* or *CAF-1* have been reported to increase the rate of spontaneous gross chromosome rearrangements (Myung et al., 2003). We observed that the reduced DLC signal in the *htbKR* mutant was significantly restored in combination with the *cac1*Δ or *asf1*Δ mutants (**Figures 3E** and **3F**). This further suggests that in the absence of H2Bubi, chromatin becomes more compacted due to the deficiency of the proteasome-dependent histone degradation pathway, leading to interference with nascent D-loop formation. Thus, reducing histone occupancy alleviates the defect.

The linker DNA serves to connect nucleosome core particles. Additionally, the complete nucleosome assembly may incorporate the linker histone H1, encoded by *HHO1* in *S. cerevisiae* (Patterton et al., 1998). This histone associates with the exterior of the core particle structure, particularly where the DNA enters and exits (**Figure 3G**). It is believed that histone H1 plays a role in organizing higher-order chromatin structure by facilitating chromatin condensation. Supporting this concept, research conducted in budding yeast has demonstrated that Hho1 restrains HR-mediated repair (Challa et al., 2021; Downs et al., 2003). We conducted an analysis involving *hho1*Δ mutants in conjunction with *htbKR* in the DLC assay. The *hho1*Δ mutants exhibited a 1.5-fold increase in nascent D-loops compared to WT cells (**Figure 3H**). The *hho1*Δ *htbKR* double mutant showed no significant difference compared to the *htbKR* mutant alone, indicating that H2Bubi likely functions upstream of Hho1 in the proteasome-dependent degradation of both core and linker histones (**Figure 3H**).

Both the INO80 and SWR complexes are recruited to site of DSBs and engage in the exchange of H2A-H2B dimers, consequently modulating the presence of the H2A variant, H2A.Z (Kalocsay et al., 2009; Lademann et al., 2017; van Attikum et al., 2007), which is encoded by *HTZ1* in *S. cerevisiae* (**Figure 3I**). To explore the impact of H2A.Z occupancy on nascent D-loop formation, we examined the DLC signal in *arp8*Δ (INO80-C), *swr1*Δ (SWR-C), and *htz1*Δ (H2A.Z) strains. Intriguingly, the *arp8*Δ mutant, characterized by elevated H2A.Z occupancy, exhibited significantly lower DLC levels compared to both the *swr1*Δ mutant, which has reduced H2A.Z occupancy, and the *htz1*Δ mutant. However, the DLC signal in both the *swr1*Δ and *htz1*Δ single mutants was significantly reduced compared to WT, suggesting that both the deposition and removal of H2A.Z are important for nascent D-loop formation (**Figure 3J**). A previous study demonstrated that H2A.Z recruits INO80-C to DSBs, where INO80-C subsequently removes H2A.Z during Rad51 filament formation (Lademann et al., 2017). Remarkably, we observed that the reduced DLC signal in the *htbKR* mutant was significantly restored in combination with the *htz1*Δ mutant (**Figure 3K**). This finding suggests that H2A.Z, INO80 and H2Bubi likely functions in the same pathway in the process of proteasome-dependent histone degradation and facilitating Rad51 filament formation.

In summary, these findings suggest that H2A.Z, INO80-C, and H2Bubi promote nascent D-loop formation through a proteasome-dependent mechanism that involves the degradation of core and linker histones and the formation of Rad51 filaments. Additionally, our data show that SWR1 promotes nascent D-loop formation, while Asf1, Cac1, and Hho1 act as negative regulators (**Figure 3L**).

### H2Bubi stabilizes extended D-loop

The regulatory influence of H2Bubi on D-loop metabolism, through nucleosome assembly and/or disassembly, could be intricate. It may impact not only the formation of nascent D-loops *via* histone degradation mechanisms but also the disassembly of nucleosome ahead of the extending D-loop and/or the stabilization of extended D-loops through nucleosome assembly (**Figure 4A**). Building upon the observation that compromising D-loop disruption factors and reducing histone occupancy can restore the nascent D-loop level to WT levels in the *htbKR* mutant (**Figures 2** and **3**), we further investigated these mutants using the DLE assay to measure the next step in HR, extension of the nascent D-loop (Piazza et al., 2018). We performed the DLE assay at a late time point, 8 hours post-DSB, as it has shown the highest level of DLE signal (**Figure 1F**) (Piazza et al., 2018).

**Figure 4.**
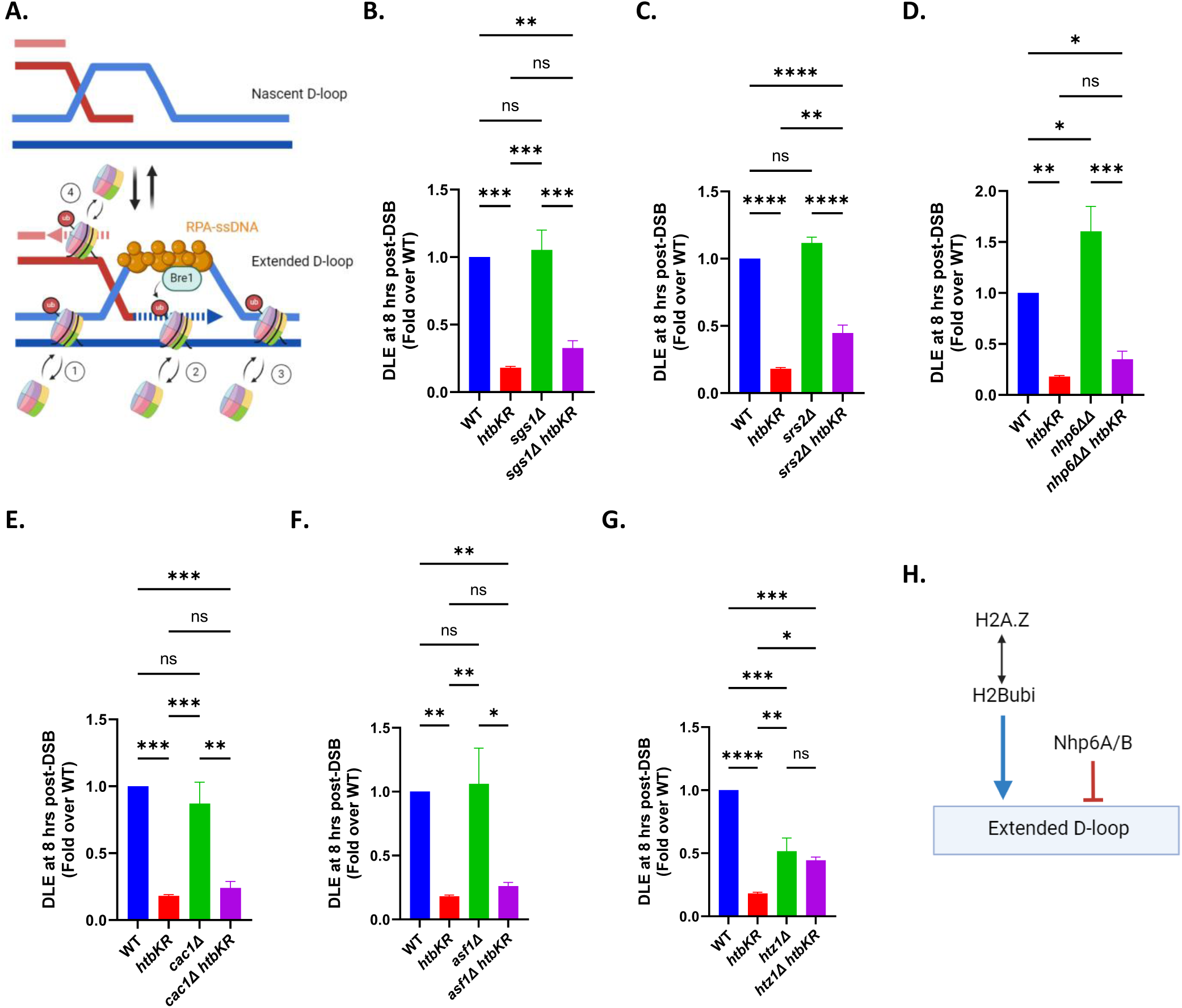
H2Bubi stabilizes extended D-loop. (**A**) An illustration depicting the process of D-loop formation and extension within the context of chromatin. H2Bubi may stabilize the extended D-loop through nucleosome assembly and/or promote nucleosome disassembly ahead of the extended D-loop. Additionally, the E3 ligase Bre1, which is responsible for H2Bubi, has been reported to interact with Rpa (Liu et al., 2021a), raising the possibility that H2Bubi is enriched at the D-loop. (**B**) Quantification of the DLE signal in WT, *htbKR*, *sgs1Δ*, and *sgs1Δ htbKR* strains. Error bars, SEM (n = 3). **: P= 0.001, ***: P< 0.0008, ns: not significant. (**C**) Quantification of the DLE signal in WT, *htbKR*, *srs2Δ* and *srs2Δ htbKR* strains. Error bars, SEM (n = 3). **: P= 0.004, ****: P< 0.0001, ns: not significant. (**D**) Quantification of the DLE signal in WT, *htbKR*, *nhp6ΔΔ* and *nhp6ΔΔ htbKR* strains. Error bars, SEM (n ≥ 2). *: P< 0.02, **: P= 0.002, ***: P< 0.0006, ns: not significant. (**E**) Quantification of the DLE signal in WT, *htbKR*, *cac1Δ* and *cac1Δ htbKR* strains. Error bars, SEM (n ≥ 2). **: P= 0.003, ***: P< 0.001, ns: not significant. (**F**) Quantification of the DLE signal in WT, *htbKR*, *asf1Δ* and *asf1Δ htbKR* strains. Error bars, SEM (n ≥ 2). *: P= 0.01, **: P< 0.004, ns: not significant. (**G**) Quantification of the DLE signal in WT, *htbKR*, *htz1Δ* and *htz1Δ htbKR* strains. Error bars, SEM (n ≥ 2). *: P= 0.01, **: P= 0.005, ***: P< 0.0006, ****: P< 0.0001, ns: not significant. (**H**) A schematic representation indicating that H2Bubi and H2A.Z promote D-loop extension, while Nhp6A/B suppress it.

We observe that the DLE signal of *sgs1*Δ, *srs2*Δ, *cac1*Δ, and *asf1*Δ single mutants exhibited no significant difference compared to WT cells at 8 hours post-DSB (**Figures 4B, 4C, 4E** and **4F**), suggesting that these mutants specifically affect the kinetic of nascent D-loop formation but have little or no effect on the kinetic of D-loop extension and by inference on the absolute D-loop levels. The *sgs1*Δ and *srs2*Δ data are consistent with previous observations (Piazza et al., 2018).

When comparing the DLE signal of *sgs1*Δ *htbKR* with *srs2*Δ *htbKR* to the *htbKR* single mutant, we observed that the DLE signal was more significantly restored in the *srs2*Δ *htbKR* mutant than in the *sgs1*Δ *htbKR* mutant (**Figures 4B** and **4C**). This observation aligns with the DLC assay results, which showed a faster kinetic of nascent D-loop formation in the *srs2*Δ *htbKR* mutant compared to the *sgs1*Δ *htbKR* mutant (**Figures 2B** and **2C**). Moreover, the faster nascent D-loop formation kinetics but limited D-loop extension of the *srs2*Δ *htbKR* double mutant suggests that H2Bubi may have an additional function in extended D-loop progression and/or stabilization. Intriguingly, the *nhp6Δ*Δ *htbKR*, *cac1*Δ *htbKR*, and *asf1*Δ *htbKR* mutants exhibited no significant difference compared to the *htbKR* single mutant by DLE assay (**Figures 4D, 4E** and **4F**). This suggests that H2Bubi may stabilize extended D-loop extension through nucleosome assembly, while its role in nascent D-loop formation involve histone degradation mechanisms. This could be mediated by recruitment of Rad6-Bre1 through its interaction with RPA which binds to the displaced strand of the D-loop (Eggler et al., 2002; Liu et al., 2021a).

The *nhp6*ΔΔ mutant exhibits a 1.5-fold increase in extended D-loop compared to WT cells (**Figure 4D**), while nascent D-loop levels remained largely unchanged (**Figure 3C**). This suggests that reducing nucleosome occupancy through *nhp6*ΔΔ promotes D-loop extension. Interestingly, the *htz1*Δ mutant displayed a 50% decrease in extended D-loop levels compared to the WT (**Figure 4G**). Note that this effect was stronger than the 30% reduction in nascent D-loop levels (**Figure 3K**), suggesting a possible function of H2A.Z in stabilizing extended D-loop. Furthermore, the *htz1*Δ *htbKR* double mutant showed no significant difference in DLE signal compared to the *htz1*Δ single mutant (**Figure 4G**), suggesting that H2A.Z functions upstream of H2Bubi for control of D-loop extension. This observation aligns with the DLC assay results, which showed a restored nascent D-loop level in the *htz1*Δ *htbKR* mutant (**Figure 3K**).

Together, these results suggest that H2A.Z and H2Bubi are epistatic in stabilizing extended D-loops, likely by finely tuning nucleosome assembly and disassembly. Additionally, our data reveal a novel function of Nhp6A/B as negative regulators of D-loop extension (**Figure 4H**).

### Excessive resection counteracts nascent D-loop formation

H2Bubi likely promotes nascent D-loop formation by facilitating histone degradation. Next, we were curious whether H2Bubi also regulates D-loop metabolism by controlling DNA end resection through the recruitment of Rad9 (Giannattasio et al., 2005; Wysocki et al., 2005). The recruitment of Rad9 is orchestrated by chromatin binding facilitated by H2A phosphorylation and Dot1-mediated H3K79 methylation, as well as Dpb11 binding through the 9-1-1 DNA checkpoint clamp (**Figure 5A**) (Waterman et al., 2020). Importantly, in cells lacking H2Bubi, Rad9 recruitment to chromatin *via* H3K79 methylation is disrupted, while H2A phosphorylation and Dpb11 binding are preserved, thereby maintaining proficiency in checkpoint activation through the 9-1-1 clamp (Giannattasio et al., 2005). This scenario provides an opportunity to analyze the specific role of Rad9 in controlling resection during D-loop metabolism through analysis of the *htbKR* mutant.

**Figure 5.**
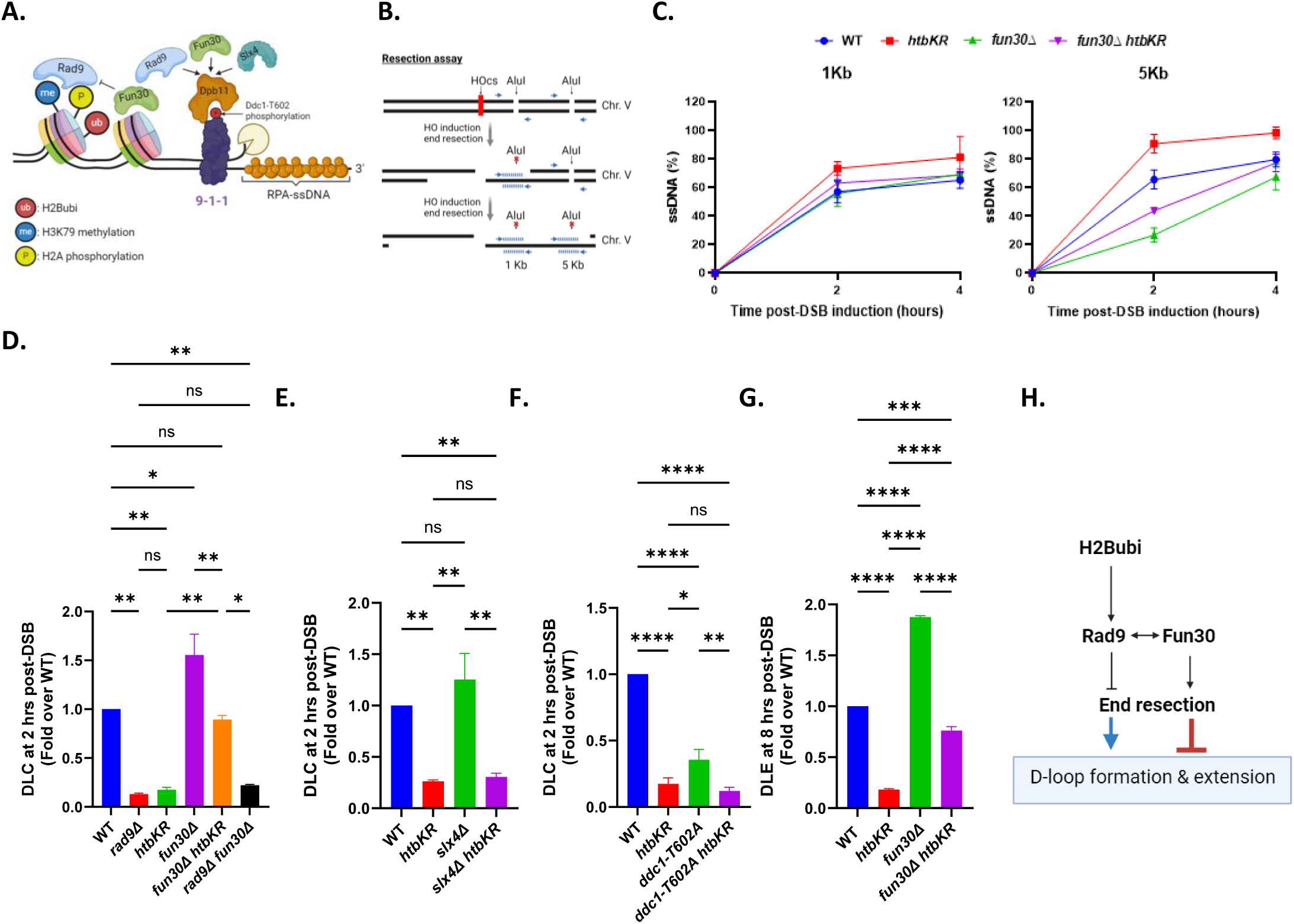
Excessive resection counteracts nascent D-loop formation and extension. (**A**) Model illustrating the role of H2Bubi in regulating DNA end resection through Rad9 recruitment. Rad9 can be recruited to chromatin *via* H3K79 methylation and H2A phosphorylation, as well as through the Dpb11 9-1-1 axis. Fun30 competes with Rad9 for chromatin binding and Dpb11 9-1-1 binding. Slx4 specifically affects Rad9 binding to the Dpb11 9-1-1 axis. (**B**) Schematic representation illustrating the quantitative PCR (qPCR) assay for monitoring DNA end resection. (**C**) Plot showing the percent of single-stranded DNA among HO cut DNA at each time point by qPCR analysis. Error bars, SEM (n = 3). (**D**) DLC signal in WT, *rad9Δ*, *htbKR*, *fun30Δ*, *fun30Δ htbKR* and *rad9Δ fun30Δ* strains. Error bars, SEM (n ≥ 2). *: P< 0.02, **: P< 0.008, ns: not significant. (**E**) DLC signal in WT, *htbKR*, *slx4Δ and slx4Δ htbKR* strains. Error bars, SEM (n ≥ 2). **: P< 0.009, ns: not significant. (**F**) DLC signal in WT, *htbKR*, *ddc1-T602A* and *ddc1-T602A htbKR* strains. Error bars, SEM (n ≥ 2). *: P= 0.01, **: P= 0.006, ****: P< 0.0001, ns: not significant. (**G**) Quantification of the DLE signal in WT, *htbKR*, *fun30Δ* and *fun30Δ htbKR* strains. Error bars, SEM (n ≥ 2). ***: P= 0.0003, ****: P< 0.0001, ns: not significant. (**H**) Schematic representation showing that H2Bubi affects nascent D-loop formation and extension by coordinating DNA end resection through Rad9 chromatin recruitment.

Fun30, a chromatin remodeler, has been identified as a competitor of Rad9 for chromatin and Dpb11 binding (Bantele et al., 2017; Chen et al., 2012; Costelloe et al., 2012; Eapen et al., 2012) (**Figure 5A**). Thus, we hypothesized that deletion of *FUN30* would lead to a deceleration of resection kinetics in the *htbKR* mutant. To test the hypothesis, we assessed resection kinetics using a well-established qPCR-based assay (Zierhut and Diffley, 2008). Primers were strategically designed to flank the *Alu*I cutting sites located 1 kb and 5 kb from the HO cut site. Since *Alu*I cannot cut ssDNA, this design allows for the amplification of PCR products from the ssDNA region generated by resection (**Figure 5B**). Indeed, we observed hyper-resection kinetics in the *htbKR* mutant and slowed resection kinetic in the *fun30*Δ mutant, consistent with earlier findings (Chen et al., 2012; Zheng et al., 2018) (**Figure 5C**). Moreover, we noted a significant reduction in resection kinetics in the *fun30*Δ *htbKR* when compared to the *htbKR* single mutant (**Figure 5C**). Control experiments using Western blots showed that that *fun30*Δ does not affect H2Bubi levels (**Figure S5A**).

To investigate whether resection kinetics impacts nascent D-loop formation, we compared the levels of nascent D-loop formation between WT, *rad9Δ, htbKR*, *fun30*Δ, *fun30*Δ *htbKR* and *rad9Δ fun30*Δ using the DLC assay at the 2 hour timepoint. Surprisingly, we observed that the *fun30*Δ strain, which exhibited slower resection kinetics, displayed increased nascent D-loop levels (**Figure 5D**). Conversely, the *rad9*Δ and *htbKR* strain, characterized by faster resection kinetics, showed reduced nascent D-loop levels at 2 hours (**Figure 5D**), suggesting an inverse relationship between resection kinetics and nascent D-loop levels. Notably, *fun30*Δ rescued the reduced nascent D-loop levels in the *htbKR* mutant but not in the *rad9*Δ mutant, indicating that Fun30 primarily competes with Rad9 in controlling end resection (**Figure 5D**). Consistent with this, a previous study found that *fun30*Δ does not slow down hyper-resection in the *rad9*Δ mutant (Chen et al., 2012). Additionally, we observed that the reduced DLC signal in the *htbKR* mutant was restored to WT levels in the *htbKR nhp6*ΔΔ double mutant (**Figure 3C**), but not in the *rad9Δ nhp6*ΔΔ double mutant (**Figure S5B**). This suggests that H2Bubi modulates chromatin structure for nascent D-loop formation, whereas Rad9 does not. Through examining Slx4, a protein known to compete with Rad9 for binding to Dpb11 but lacking influence on Rad9 chromatin binding (Dibitetto et al., 2016; Ohouo et al., 2013), we found that the *slx4*Δ mutation does not rescue the decreased nascent D-loop levels in the *htbKR* mutant (**Figure 5E**). This finding lends support to the hypothesis that H2Bubi primarily impacts Rad9 chromatin binding rather than Dpb11 binding. To further support this notion, we disrupted the Mec1-dependent Ddc1 phosphorylation site (*ddc1-T602R*), which is crucial for the recruitment of Rad9, Slx4, and Fun30 to Dpb11 (Bantele et al., 2017; Ohouo et al., 2013; Pfander and Diffley, 2011; Puddu et al., 2008). This mutation failed to rescue the DLC defect observed in the *htbKR* mutant, indicating that H2Bubi operates independently of the Dpb11 9-1-1 axis in nascent D-loop formation (**Figure 5F**).

Building on the observation that slowing down resection through *fun30*Δ can restore nascent D-loop levels to WT levels in the *htbKR* mutant (**Figure 5D**), we further investigated these mutants using the DLE assay to measure the next step in HR: the extension of the nascent D-loop. The *fun30*Δ mutant exhibits a 2-fold increase in extended D-loop levels compared to WT cells (**Figure 5G**), a more pronounced effect than the 1.5-fold increase observed in nascent D-loop levels (**Figure 5D**). This suggests a novel role for Fun30 in negatively regulating D-loop extension. Additionally, the DLE signal was partially restored in the *fun30*Δ *htbKR* double mutant compared to the single mutants *htbKR* and *fun30*Δ (**Figure 5G**). This finding aligns with the DLC assay results, which showed restored nascent D-loop levels in the *fun30*Δ *htbKR* mutant (**Figure 5D**), indicating that Fun30 acts epistatically to H2Bubi in controlling D-loop formation and extension.

Taken together, these data suggest that excessive resection negatively feeds back on nascent D-loop formation and that H2Bubi plays a role in this process likely through regulating Rad9 chromatin binding (**Figure 5H**).

### H2Bubi affects DSB repair pathway usage and outcome

To explore the impact of H2Bubi on DSB repair pathway usage and outcomes, we used a previously described DSB-induced mitotic recombination assay. This assay monitors both reciprocal and nonreciprocal exchange end products in diploid cells monitoring crossover (CO) and non-crossover (NCO) outcomes as well as BIR and chromosome loss along with additional events leading to loss of heterozygosity (LOH) (Ho et al., 2010). In this system, an *I-Sce1* cut site is introduced within the *ADE2* locus (*ade2-I*) on one chromosome XV (Chr. XV), while the other homolog carries an inactive allele of *ADE2* (*ade2-n*). The repair outcome is assessed using two antibiotic resistance genes, *HPH* and *NAT*. Additionally, *MET22* and *URA3* were employed to trace chromosome loss events (**Figure 6A**).

**Figure 6.**
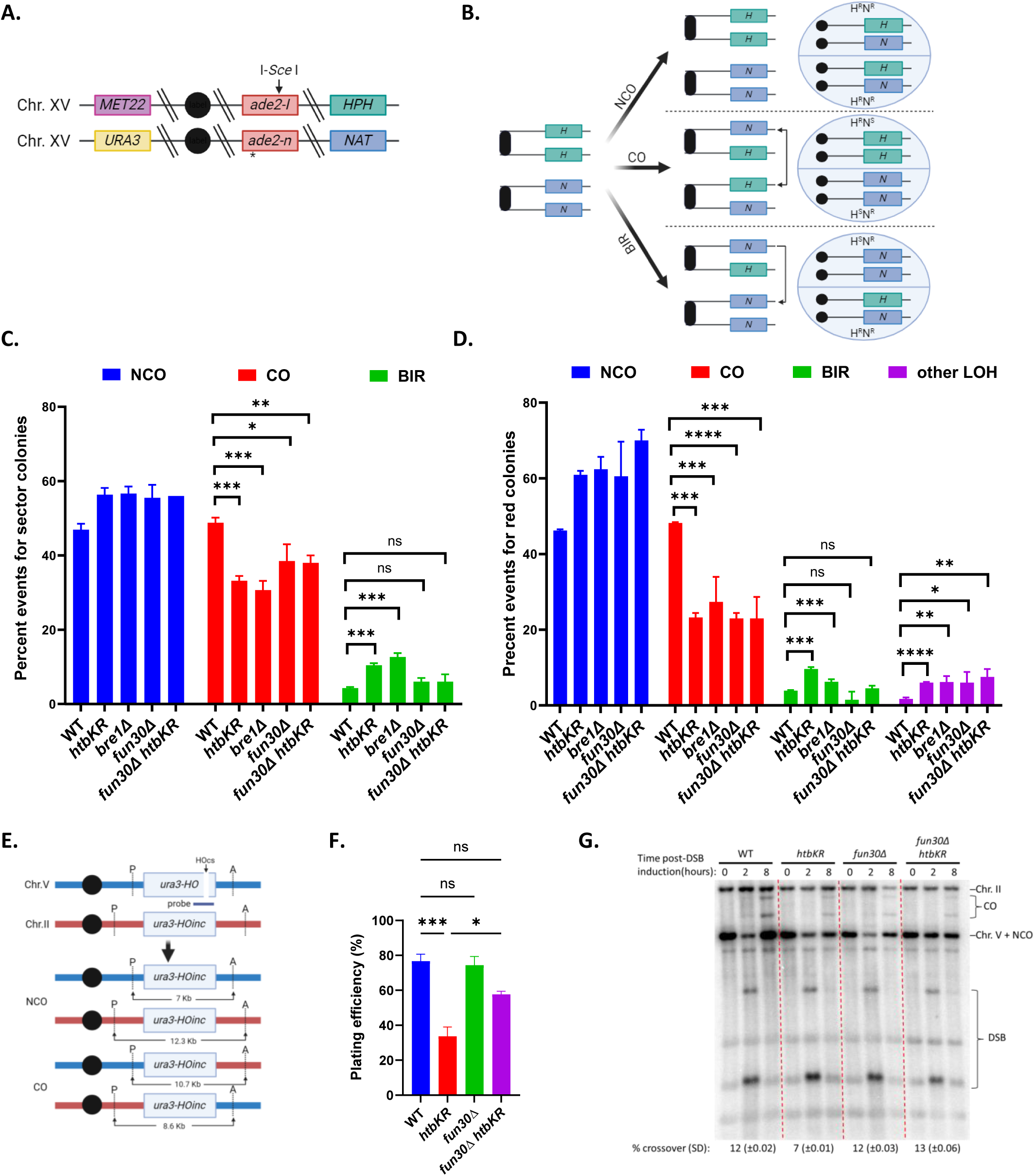
H2Bubi affects DSB repair pathway usage and outcome. (**A**) Schematic representation of the diploid strain, indicating the *I-SCE*1 cut site and the selection markers for determining repair outcomes and monitoring chromosome loss events (See text for details). (**B**) Schematic representation of repair outcomes, illustrating unchanged Nat and Hph markers in non-crossover (NCO) repair, reciprocal exchange of Nat and Hph markers in crossover (CO) repair, and non-reciprocal exchange of Nat and Hph markers in break-induced replication, resulting in homozygosis of the Nat marker. H: Hph, N: Nat, R: resistance, S: sensitive. (**C**) Percentage of non-crossover (NCO), crossover (CO), and break-induced replication (BIR) events for sector colonies among the indicated strains. Error bars, SEM (n ≥ 2). ***:P<0.0007. **:P< 0.01. *:P<0.04. ns: not significant. (**D**) Percentage of occurrences for each strain within the red colonies. Error bars, SEM (n ≥ 2). ****:P<0.0001, ***:P<0.0008, **:P< 0.003, *:P<0.03. ns: not significant. (**E**) Schematic representation of the chromosome II-chromosome V ectopic recombination assay, depicting the distance between the HO endonuclease cut site (HO) and the heterology *ApaL*I (A) and *Pvu*II (P) sites located outside the region of shared homology were utilized to detect crossover (CO). Sizes of parental, non-crossover (NCO), and CO fragments are indicated. (**F**) Plating efficiency was assessed by counting colony formation on YP-Gal plates divided by YP-Glu plates for each of the indicated strains across three independent trials. Error bars, SEM (n = 3). *: P= 0.02, ***: P< 0.0007, ns: not significant. (**G**) Southern blot analysis was conducted on DNA extracted from cells sampled at 0, 2, and 8 hours post double-strand break (DSB) induction. Genomic DNA from each strain was digested with *Pvu*II and *ApaL*I enzymes, then probed with a fragment of Chromosome V carrying the *URA3* gene. The percentage of crossover product at the 8-hour time point was calculated using densitometer quantification. This percentage was obtained by dividing the intensity of crossover bands by the total DNA content (Chromosome II, Chromosome V, and crossover). Subsequently, the crossover (CO) level was normalized to the plating efficiency obtained from (E). The numbers in parentheses represent the standard deviation (SD) values obtained from three independent trials.

Following *GAL-I-Sce1* induction, DSBs at *ade2-I* were repaired *via* gene conversion with *ade2-n* when both sister chromatids were cleaved in G2 cells. The pattern of gene conversion tracts was determined by colony color: solid white for short tract gene conversion on both sisters, sectored for short tract on one and long tract on the other, and solid red for long tract on both sisters (**Figure S6A**). We found a similar ratio of long tract *versus* short tract gene conversion (**Figure S6B**) and observed no chromosome loss events in any of the mutants tested (**Table S1**).

The repair outcomes were determined by the sensitivity of cells to the antibiotic drugs Nourseothricin (Nat) and Hygromycin B (Hph) (**Figure 6B**). Repair to NCO events results in Hph and Nat resistance (H^R^N^R^/H^R^N^R^), while repair to CO events leads to either Hph or Nat sensitivity (H^R^N^S^/H^S^N^R^). Repair by BIR events result in Nat resistance (H^S^N^R^/H^R^N^R^) (**Figure 6B**). The most definitive determinant of repair outcomes was observed through the analysis of sectored colonies. In WT cells, sectored colonies exhibited a nearly even distribution of CO to NCO events, along with a small percentage of BIR events, consistent with the original study (**Figure 6C**) (Ho et al., 2010). We observed a reduction in CO events in *bre1*Δ and *htbKR* mutants, accompanied by an increase in BIR events, but only in *bre1*Δ and *htbKR* mutants; no such increase was observed in *fun30*Δ and *fun30*Δ *htbKR* mutants (**Figure 6C**). The reduced CO was also evident in *bre1*Δ and *htbKR* mutants when measuring solid red colonies (**Figure 6D**) and solid white colonies (**Figure S6C**). Similarly, the increased BIR was also observed in *bre1*Δ and *htbKR* mutants when analyzing solid red colonies (**Figure 6D**), whereas no such increase was noted in solid white colonies. Solid white colonies underwent two short tract conversion events, for which NCO is the preferred repair outcome. In these cells, only a limited amount of CO and BIR were observed even in WT (**Figure S6C**). Remarkably, the occurrence of BIR events in *fun30*Δ *htbKR* mutants closely resembled that of the WT cells across all types of colonies examined (sectored, red, and white) (**Figures 6C, 6D** and **S6C**). In sum, these data show that the influence of H2Bubi on D-loop metabolism can significantly affect HR sub-pathway usage and outcome.

To validate the observed repair outcome in the diploid system, we employed a well-established physical recombination assay in haploid cells (Aylon et al., 2003). Haploid strains featuring an HO cut site insertion within the native *URA3* locus (*ura3-HO*) on chromosome V (Chr. V), alongside a 5.6-kb *ura3-HOinc* (non-cleavable) fragment integrated at the *LYS2* locus on chromosome II (Chr. II) (**Figure 6E**). Upon *GAL-HO* induction, the DSB on Chr. V undergoes repair *via* gene conversion, transferring the *ura3-HOinc* allele from Chr. II to Chr. V. To assess DSB repair efficiency, we examined the plating efficiency (PE) of each strain on galactose-containing medium compared to glucose-containing medium. Our analysis revealed a significantly lower PE for the *htbKR* mutant compared to the WT (**Figure 6F**), demonstrating an involvement of H2Bubi in DSB repair. To ascertain the repair outcome, we induced HO in liquid cultures and isolated DNA at various time points post-HO induction for restriction digestion and Southern blot hybridization. Mutants displaying decreased viability may be associated with the use of the BIR pathway, leading to the loss of essential genes downstream of the *ura3-HO* site, thereby resulting in lower PE. Since BIR and CO events are indistinguishable in the physical assay, we corrected the CO signal determined by Southern blot hybridization to the PE of all examined strains (**Figure 6G**), as done in previous studies (Mazon et al., 2012; Mazon and Symington, 2013b). The analysis revealed a 2-fold decrease in CO events in the *htbKR* mutant compared to the WT (**Figure 6G**). Intriguingly, *fun30*Δ rescues both PE and CO in the *htbKR* mutant, highlighting the inverse relationship between H2Bubi and Fun30 in DSB repair (**Figures 6F** and **6G**).

Collectively, we conclude that the impact of H2Bubi on D-loop metabolism can influence the usage of HR sub-pathways. If not tightly regulated, BIR may be used for repair, a process demonstrated to be mutagenic (Kockler et al., 2021).

## DISCUSSION

H2B mono-ubiquitylation (H2Bubi) has previously been associated with DSB repair (Challa et al., 2021; Hung et al., 2017; Moyal et al., 2011; Nakamura et al., 2011; Zheng et al., 2018), yet the precise molecular mechanisms underlying its involvement in D-loop metabolism and its impact on repair outcome have remained poorly understood. In this study, we use the novel and validated DLC and DLE assays to identify and measure critical recombination intermediates, namely nascent and extended D-loops (Piazza et al., 2018; Piazza et al., 2019). These assays offer a unique advantage with their ability to directly detect these intermediates, setting them apart from techniques that rely on measuring the recruitment of recombinase Rad51 and physical or genetic repair endpoints. Rad51 measurement methods can suffer from non-specific issues, as Rad51 presence can be triggered by various forms of DNA damage beyond HR, for example stalled forks for fork protection (Mason et al., 2019). Furthermore, the DLC and DLE assays diverge from genetic analysis, which primarily focuses on providing insights into repair outcomes but lacks the necessary detail to fully understand molecular mechanism of the recombination process.

By using DLC and DLE assays to monitor D-loop metabolism, alongside physical and genetic assays to detect repair outcomes in cells lacking H2Bubi and various mutants affecting chromatin, we have uncovered multi-faceted functions of H2Bubi in D-loop metabolism. H2Bubi finely regulates chromatin structure: Firstly, it facilitates histone degradation, thus promoting the formation of nascent D-loops. Secondly, it potentially stabilizes extended D-loops likely through nucleosome assembly. Thirdly, H2Bubi regulates DNA resection: It coordinates the kinetics of DNA resection with the kinetics of D-loop formation and extension by recruiting Rad9 (**Figure 7**).

**Figure 7.**
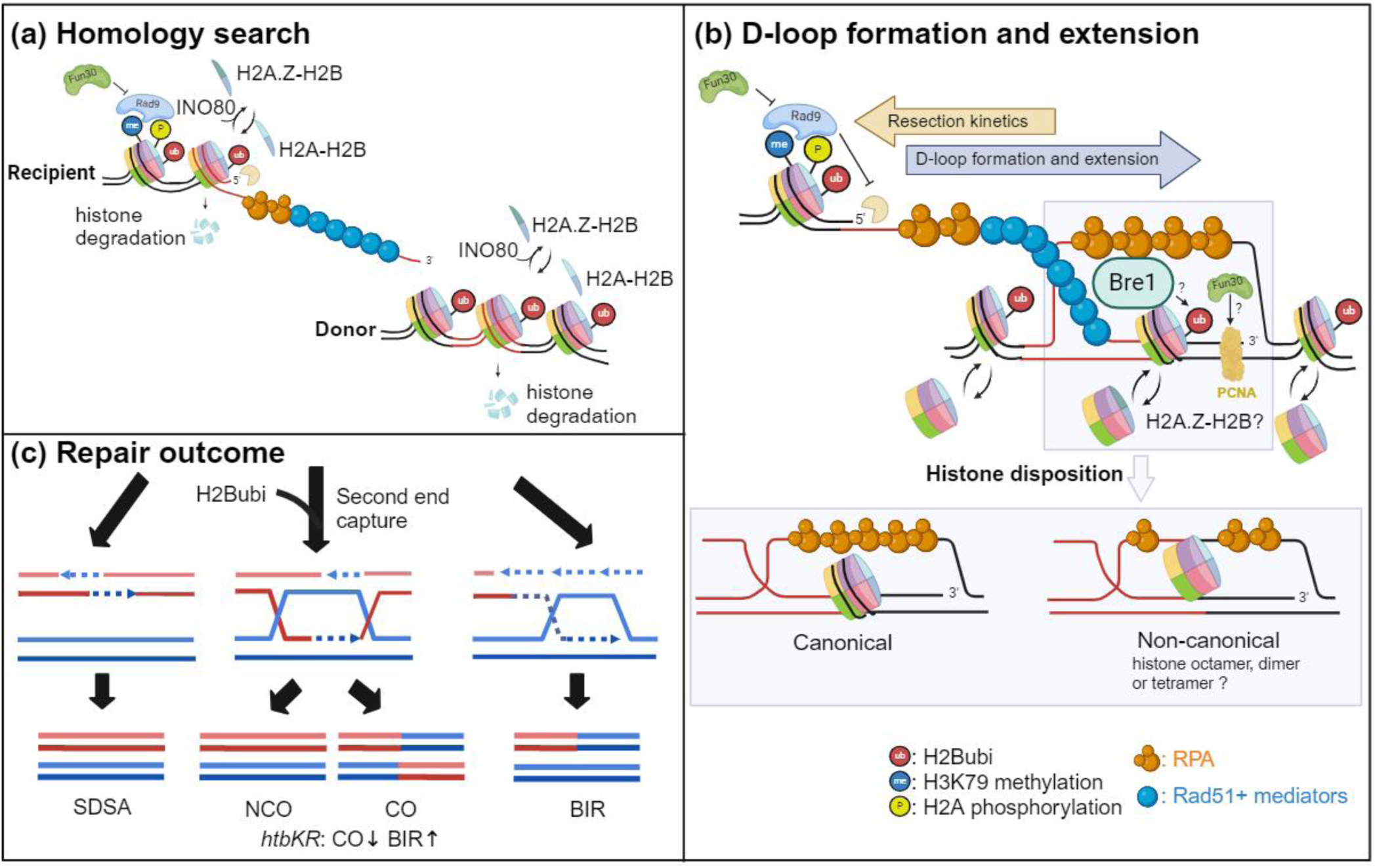
Model depicting the regulation of D-loop metabolism and repair outcomes by H2Bubi. H2Bubi may regulate D-loop metabolism through several mechanisms: (**A**) by promoting histone degradation to facilitate homology search and nascent D-loop formation; (**B**) by coordinating DNA resection with D-loop formation and extension through Rad9 recruitment and stabilizing the extended D-loop via nucleosome assembly. The epistatic relationship between H2Bubi, Fun30, and H2A.Z suggests potential cooperation in DNA resection and D-loop metabolism; (**C**) H2Bubi may also influence second-end capture, potentially shifting repair from crossover (CO) to break-induced replication (BIR).

### H2Bubi facilitates nascent D-loop formation

Our study offers the first evidence demonstrating the impact of H2Bubi on nascent D-loop formation and extension. Notably, the deficiency in nascent D-loop formation can be restored by eliminating factors involved in the D-loop disruption pathways identified in our previous research (Piazza et al., 2019), underscoring the critical role of the D-loop disruption pathway in monitoring nascent D-loops. Furthermore, manipulating chromatin structure by removing factors known to alter chromatin structure through various mechanisms influences nascent D-loop levels to varying degrees. Our studies provide several line of evidences to suggest that H2bubi fine-tunes chromatin structure for HR: First, we demonstrated that *ARP8* deletion, previously associated with compacted chromatin, resulted in a dramatic reduction in nascent D-loop formation, similar to the *htbKR* mutant, which is congruent with previous studies suggesting that Rad6 and Bre1 recruitment upon DNA damage depends on INO80 (Challa et al., 2021). Conversely, the *NHP6A/B* deletion, previously linked to expanded chromatin, showed complete rescue of the D-loop defect in *htbKR*, likely due to a 20% reduction in histone occupancy (Celona et al., 2011; Challa et al., 2021; Cheblal et al., 2020). Moreover, our epistasis analysis revealed that Hho1 acts downstream of H2Bubi, consistent with previous studies proposing core and linker histone degradation upon DNA damage (Challa et al., 2021; Hauer et al., 2017). Our study further defines H2Bubi’s role as upstream of linker histone Hho1. Second, our epistasis analysis shows that the histone chaperones Asf1 and CAF-1 act upstream of H2Bubi, consistent with their role in promoting the assembly and disassembly of nucleosomes. Our results demonstrate that reducing histone occupancy by deleting *ASF1* or *CAC1* reverses the decompaction defect observed in the *htbKR* mutant. Third, our epistasis analysis reveals that the histone variant H2A.Z acts upstream of H2Bubi, which is intriguing. One possible explanation is that in the absence of H2Bubi, H2A.Z inhibits Rad51 filament formation, similar to previous reports showing restoration of presynaptic filament formation and HR in INO80-C-deficient mutants when H2A.Z is absent (Lademann et al., 2017). It is worth noting that the restored nascent D-loop level in the *htz1*Δ *htbKR* double mutant can be explained by the restored presynaptic filament formation. Furthermore, we observed that the deposition of H2A.Z also play a role in promoting nascent D-loop formation, as evidenced by the reduced DLC levels in *swr1*Δ and *htz1*Δ mutants. Our results reveal a complex role for H2A.Z in nascent D-loop formation, consistent with previous studies demonstrating that H2A.Z is deposited to promote DNA resection and later removed during Rad51 filament formation (Kalocsay et al., 2009; Lademann et al., 2017; van Attikum et al., 2007). In conclusion, we infer that INO80 acts upstream of H2Bubi through H2A.Z removal to promote proteasome-dependent histone degradation, which promotes Rad51 filament formation as a prerequisite for homology search and nascent D-loop formation (**Figure 7**).

### H2Bubi promotes nascent D-loop formation by facilitating histone degradation and stabilizes extended D-loop *via* nucleosome assembly

H2Bubi stands out as one of the most significant chromatin marks known in the regulation of chromatin structure and compaction (Fierz et al., 2011). However, the mechanism by which H2Bubi fine-tunes chromatin structure to facilitate HR during D-loop metabolism remains unclear. Our study provides the first evidence demonstrating that H2Bubi promotes nascent D-loop formation through histone degradation (**Figure 3**), while stabilizing the extended D-loop *via* chromatin assembly (**Figure 4**). This hypothesis is supported by several lines of evidence: First, removing D-loop disruption pathways significantly recovered nascent D-loop levels in the *htbKR* mutant, while extended D-loop levels remained low (**Figure S7A, D-loop disruption category**). Second, the reduction of histone occupancy restored nascent D-loop levels but did not rescue extended D-loop levels in the *htbKR* mutant. This suggests that H2Bubi stabilizes the extended D-loop through chromatin assembly, rather than promoting chromatin disassembly ahead of it (**Figure S7A, chromatin structure category**). This may resemble DNA replication, where H2Bubi promotes chromatin assembly behind the replication fork (Lin et al., 2014; Trujillo and Osley, 2012). Similarly, we speculate that H2Bubi may stabilize the extended D-loop by chromatin assembly at or behind the D-loop (**Figure 7**). This may involve not only full nucleosomes but possibly also histoner dimer or tetramers and potentially non-canonicaly ways of engaging with DNA as speculated in (**Figure 7B**).

Moreover, the direct interaction between Bre1 with RPA, Rad51 and Srs2 (Liu et al., 2023; Liu et al., 2021a) raises the possibility that Bre1 may be recruited to extended D-loops through RPA, Rad51, and Srs2 binding, enhancing local H2Bubi levels to facilitate DNA synthesis during D-loop extension. However, this needs further investigation.

Our study observed no significant effect of Asf1 and CAF-1 on D-loop extension during the repair of a two-sided (frank) DSB. In a different functional context, recombination-dependent replication in fission yeast, both were founds required for DNA synthesis (Hardy et al., 2019; Pietrobon et al., 2014), which may reflect the difference between HR during template switching at stalled forks where HR intermediates may be stabilized by histone deposition.

Additionally, it has been shown that the human homolog of Fun30, SMARCAD1, directly interacts with the PCNA replication clamp and is preferentially enriched at unperturbed replication forks, suggesting a role in DNA replication (Lo et al., 2021). Our study reveals a novel function of Fun30 that inversely correlates with H2Bubi in D-loop extension. Whether this correlation relates to its replication function requires further investigation.

Recent cryo-electron microscopic structural studies have revealed that H2A.Z destabilizes the nucleosome structure by enhancing DNA mobility near the DNA entry/exit site, while also facilitating the formation of more regular and compact chromatin fiber in nucleosome arrays (Lewis et al., 2021). The interplay between H2Bubi and H2A.Z contributes to transcriptional regulation in a context-dependent manner (Segala et al., 2016; Wojcik et al., 2018). This suggests that H2Bubi may influence nascent D-loop formation and extension by stabilizing H2A.Z in nucleosomes or hindering its eviction, and *vice versa*.

Due to the structural similarity between the Fun30 with the INO80 and SWR complexes, Fun30 may play a role in histone dimer exchange (Awad et al., 2010; Flaus et al., 2006). It has been shown that *FUN30* genetically interacts with subunits of the SWR complex and histone H2A.Z (Krogan et al., 2003) and *FUN30* deletion results in defects in genome-wide histone variant H2A.Z occupancy, further supporting this notion (Durand-Dubief et al., 2012). Our study reveals that the crosstalk between H2Bubi, H2A.Z, INO80, SWR1, and Fun30 provides a possibility that their interaction may fine-tune chromatin structure for D-loop metabolism (**Figure 7**).

### Resection influences D-loop metabolism

The rate of resection, necessary to produce single-stranded DNA for homology search, may serve as a time-limiting factor for the homology search process. Initially, the formed 3’ ssDNA is coated by RPA, followed by Rad51 filament formation, which facilitates the search for a homologous sequence. If successful, this search is succeeded by the invasion of the template sequence. Surprisingly, hyper-resection in cells lacking H2Bubi does not result in successful strand invasion, indicating that excessive resection negatively affects D-loop levels (**Figure 5**). Consistently, we observed that slowing down resection kinetics by deleting *FUN30* restored the kinetics of nascent D-loop formation and extension in the *htbKR* mutant (**Figures 5D** and **5G**). Furthermore, we found that the reduced cell viability, decreased CO formation and increased BIR observed in the *htbKR* mutant were restored to WT levels in the *fun30*Δ *htbKR* double mutant (**Figures 6C, 6D, 6F** and **6G**).

This suggests that H2Bubi may play an additional role in sustaining the kinetics of D-loop formation and extension by limiting excessive DNA end resection through recruiting Rad9 (**Figure S7A, chromatin factors affecting Rad9 recruitment**). This finding is supported by research suggesting that Rad9 has an additional function independent of Rad53 checkpoint signaling in stabilizing D-loops by limiting the recruitment of proteins involved in D-loop disruption pathways (Ferrari et al., 2020). Additionally, current research suggests that in the absence of Rad9, hyperactivating of Mec1 signaling regulate the STR-dependent D-loop disruption pathway to influence HR outcomes and suppress gross chromosomal rearrangements (Sanford et al., 2021; Xie et al., 2024). Moreover, Rad9’s function in Rad53 checkpoint signaling has been demonstrated to prevent aberrant processing of BIR intermediates (Vasan et al., 2014). In sum, these results suggest the existence of a negative feedback loop between resection and D-loop reversal that safeguards the homology search process to limit non-allelic HR and gross chromosomal rearrangements (**Figure 7B**).

### H2Bubi affects DSB repair pathway usage and outcome

D-loops serve as a central intermediate of HR-mediated repair. Therefore, if any issues arise during D-loop metabolism, the frequencies BIR, CO, and NCO may be affected. To this end, we examined the competition between DSB repair sub-pathways using genetic and physical assays (**Figure 6**). In the DLC/E and BIR assays, DSBs are exclusively repaired by the BIR pathway (**Figure 1**) not allowing to study competition between HR sub-pathways. In competition assays, we consistently observed an increase in BIR and a decrease in CO outcomes by analysis of genetic and physical endpoints in cells lacking H2Bubi. BIR events could arise due to the failure of second end capture thus discourage the formation of double Holliday junctions and subsequent COs (Piazza and Heyer, 2019; Wright et al., 2018).

Our results provide evidence to explain why CO decreases and BIR increases in the absence of H2Bubi. The kinetics of nascent and extended D-loops are dramatically delayed in cells lacking H2Bubi, consistent with the hypothesis that limiting the lifetime of nascent and extended D-loops discourages the formation of dHJs and subsequent crossovers. Moreover, our observation is supported by a previous study demonstrating reduced meiotic crossovers upon depletion of the E3 ligase RNF20 for H2Bubi in mice (Xu et al., 2016). Finally, inappropriate DNA resection may interfere with engaging the second end either by annealing and strand invasion.

We conclude that the effect of H2Bubi on D-loop metabolism can influence repair pathway usage (**Figure 7C**). Specifically, H2Bubi plays a crucial role in preventing break-induced replication, a sub-pathway of DNA double-strand break repair known to culminate in genomic instability.

### Limitations of the study

In this study, we reveal that H2Bubi plays multi-faceted roles in regulating the kinetics of nascent D-loop formation and extension. However, due to the technical constraints of visualizing nucleosomes on extended D-loops, we still do not fully understand how H2Bubi stabilizes extended D-loops. Additionally, the molecular mechanism by which H2Bubi coordinates DNA resection and D-loop formation is lacking. Further studies analyzing resection intermediates would provide a clearer interpretation of this molecular mechanism. H2Bubi controls H3 methylation through Dot1-mediated H3K79 and Set1-mediated H3K4 methylation (Game and Chernikova, 2009). While previous studies suggest that the HR repair function of H2Bubi is independent of H3K4 or H3K79 methylation (Zheng et al., 2018), we cannot definitively state whether the effect we observe are directly or indirectly caused by H2Bubi.

## MATERIAL AND METHODS

### Yeast Strains and Growth Conditions

The yeast strains and their genotypes (W303 *RAD5* background) utilized in this study are listed in (**Table S2**). Gene disruptions were achieved by replacing the indicated gene using a PCR-based strategy (Guldener et al., 1996). All yeast cells were cultured in YP medium supplemented with 2% (vol/vol) dextrose (1% yeast extract, 2% peptone). For DLC/DLE assays, cells were initially grown in YPD medium (1% yeast extract, 2% peptone, 2% dextrose), then diluted and switched to YEP-lactate medium (1% yeast extract, 2% peptone, 2% lactate). For haploid endpoint assays, cells were grown in YEP-lactate medium before adding 2% galactose for *GAL-HO* induction. For diploid endpoint assays, cells were cultured in YPR medium (1% yeast extract, 2% peptone, 2% raffinose) before adding 2% galactose for *GAL-ISCEI* induction. All analyses were conducted during the log phase of growth at 30 °C.

### D-loop capture and extension assays

For D-loop capture experiments, all strains were in the W303 *RAD5* background. They contain a copy of the GAL1/10 driven HO endonuclease gene at the *TRP1* locus on chr. IV. A point mutation inactivates the HO cut-site at the mating-type locus (MAT) on chr. III (*MATa-inc*). The DSB-inducible construct contains the 117 bp HO cut-site, a 2,086 bp-long homology A sequence (+4 to +2090 of the LYS2 gene), and a 327 bp fragment of the PhiX174 genome flanked by multiple restriction sites. The D-loop capture and extension assays were conducted following established protocols with slight adjustments (Piazza et al., 2018; Piazza et al., 2019; Reitz et al., 2022). Specifically, zymolyase-lysed cells were immediately subjected to restriction digestion, ligation, and DNA purification steps following hybridization with oligonucleotides. The control experiments that monitoring DSB formation, ligation efficiency, and a normalization locus are shown (**Figures S1** to **S5**).

### Determination of BIR frequency

BIR frequency determination followed the method outlined in (Donnianni and Symington, 2013). In brief, cells were cultivated to exponential phase in YEP-lactate medium and then plated on YPD plates. After 3 days, colonies were counted and subsequently replicated onto synthetic complete medium lacking lysine (*LYS2* drop-out) plates or YPD plates containing geneticin (G418). Cell viability post-HO induction was calculated by dividing the number of colonies on YP galactose plates by those on YP glucose plates. The percentage of cells repairing *via* BIR was determined by dividing the number of cells on *LYS2* drop-out plates by the number on YP galactose plates, normalized to the number on YPD. BIR frequencies were determined three times for each strain.

### Detection of BIR products formation

Cells were cultured in YEP-lactate medium, followed by the addition of 2% galactose to induce HO endonuclease expression. Genomic DNA (25 ng) was then subjected to PCR amplification using Phusion High-Fidelity DNA Polymerase with the following cycling conditions: initial denaturation at 98°C for 30 seconds, followed by 25 cycles of denaturation at 98°C for 10 seconds and annealing/extension at 72°C for 150 seconds in a 25 μl reaction volume. BIR product detection utilized P1 and P2 primers, while HO cut detection employed D1 and D2 primers. Normalization of P1-P2 and D1-D2 products was achieved using C1 and C2 primers, as described previously (Donnianni and Symington, 2013).

### DNA end resection determined by qPCR-based assay

Resection assay was performed as described in (Zierhut and Diffley, 2008). Cells were cultured in YEP-lactate medium and induced with 2% galactose. Genomic DNA (150 ng) was collected at 0, 2, and 4-hour time points and digested with 10 units of *Alu*I at 37°C for 3 hours. Subsequently, the digested DNA samples or mock digests were diluted in ice-cold ddH2O and kept on ice for quantitative PCR (qPCR) analysis. Primers flanking the *Alu*I cutting site and located 1 and 5 kb away from the HO cut site were used to quantify end resection. A control primer pair targeting a region on *ARG4* lacking the *Alu*I site was used for normalization. qPCR was conducted using Light Cycler 480 SYBR Green I Master with the Light Cycler 480 System (Roche), following the program: 95°C for 15 s, 61°C for 12 s, 72°C for 15 s, repeated for 50 cycles. Triplicate reactions were performed for all primer pairs, and the average threshold cycle value was calculated for each sample. The percentage of ssDNA present at each time point was determined using the formula described in (Zierhut and Diffley, 2008).

### Recombination outcome determined by haploid endpoint assay

The haploid yeast strain utilized for the ectopic recombination assay was previously detailed (Aylon et al., 2003). Plating efficiency was calculated by dividing the count of colonies from YP galactose plates by those from YP glucose plates. The distribution of crossover (CO) products was assessed *via* Southern blot hybridization, employing a *URA3* probe on *ApaL*I-*Pvu*II digested genomic DNA, as outlined in (Mazon and Symington, 2013a).

### Recombination outcome determined by diploid endpoint assay

The diploid yeast strain utilized for the ectopic recombination assay was previously described (Ho et al., 2010). Cells were cultured in YPR medium until reaching the logarithmic phase, following which 2% galactose was added to induce *I-Sce*1 expression for 1.5 to 3 hours, depending on the growth rate of the wild type and each mutant. Subsequently, cells were plated on YPD and allowed to grow for two days. YPD plates were then replica plated onto YPD with hygromycin B (200 μg/ml), YPD with nourseothricin (67 μg/ml, clonNat), and SC (-Ura/-Met) plates to determine recombination outcomes and ensure proper chromosome segregation, respectively. Notably, the YPD plates were also replica plated onto re-induction (SC-Ade medium containing 2% raffinose and 1% galactose (SCR-Ade+Gal)) plates to confirm that red colonies and the red halves of sector colonies were genuine long-tract gene conversions. Any red colonies that turned white after replica plating onto re-induction plates were not counted, as this indicated that not both DSB sites were cut during galactose induction in liquid culture. Statistical significance for the number of recombination events between given strains was calculated using Student’s t-test. Independent inductions were performed at least two times for each strain.

### Western blot

Yeast cell lysates were prepared using the TCA method. Briefly, equivalent numbers of cells (4 × 10^7^) were collected and resuspended in 500 μl of ddH2O. Then, 75 μl of ice-cold NaOH/βME solution (1.85 M/7.5% final) was added, followed by vortexing and incubation on ice for 15 minutes. After adding 75 μl of ice-cold 55% TCA and another round of vortexing, the lysates were incubated on ice for 10 minutes. The supernatant was completely removed after the final wash. Samples were suspended in 30 μl of HU (high urea) buffer [8 M urea, 200 mM Tris·HCl (pH 6.8), 1 mM EDTA, 5% (wt/vol) SDS, 0.1% (wt/vol) bromophenol blue, 1.5% (wt/vol) DTT] and denatured at 60 °C for 10 minutes. Before analysis by SDS/PAGE, samples were briefly centrifuged. The anti-H2B (active motif) antibody was utilized to detect histone H2B and H2Bubi.

### Statistical analysis

The statistical analysis of DLC and DLE signals for each mutant was compared to their respective paired mutants using ordinary one-way ANOVA in Prism 10 (GraphPad Software). Additionally, the statistical analysis of genetic recombination outcomes for each mutant was compared to the wild type and was conducted using unpaired Student’s t-test.

## ACKNOWLEDGEMENTS

We express our gratitude to the members of the Heyer laboratory, especially Diedre Reitz, for their valuable discussions. We thank Susan Gasser and Marcus Smolka for insightful comments on the manuscript. We also extend our thanks to Lorraine Symington, Marcus Smolka, and Cheng-Fu Kao for providing yeast strains and plasmids. This work received support from NIH awards R01GM58015 and R01GM137751 granted to W.D.H., and the Cancer Center Core Support grant P30CA093373. Additionally, S.H.H acknowledges partial support from a fellowship provided by Academia Sinica, Taiwan. All figures illustrating the models were created with BioRender.com.

## AUTHOR CONTRIBUTIONS

S.H.H. and W.D.H. conceived the project and drafted the manuscript. S.H.H. designed, conducted, and analyzed the experiments. Y.L. contributed to the experimental work and analysis.

## DECLARATION OF INTERESTS

The authors declare no conflict of interests.

---

**Figure S1. Control experiments for the DLC and DLE assays of WT, *bre1*Δ and *htbKR* strain. Related to figure 1.**

The DNA loading control experiment measures the amplification cycle (Cp value) of the control locus, *ARG4*. The efficiency of double-strand break (DSB) induction is determined by comparing the percent amplification from the HO cut site at 2 hours to that at 0 hours. Additionally, the intramolecular ligation efficiency assesses the ligation efficiency of a circularized DNA fragment from Chr.VIII following *EcoR*1 digestion for DLC assay control, and a circularized DNA from Chr.XII following *Hind*III digestion for DLE assay control (Reitz et al., 2022). (**A**) Plot of individual values of the HO cutting efficiency of the DLC assay of each strain. (**B**) Plot of individual values of the DNA loading control of the DLC assay of each strain. (**C**) Plot of individual values of the intramolecular ligation control of the DLC assay of each strain. (**D**) Plot of individual values of the DNA loading control of the DLE assay of each strain. (**E**) Plot of individual values of the intramolecular ligation control of the DLC assay of each strain.

**Figure S2. Control experiments for the DLC assay of *htbKR*, as well as for genes that implicated in D-loop disruption, examined individually or in double mutant combinations. Related to figure 2.**

The DNA loading control experiment measures the amplification cycle (Cp value) of the control locus, *ARG4*. The efficiency of double-strand break (DSB) induction is determined by comparing the percent amplification from the HO cut site at 2 hours to that at 0 hours. Additionally, the intramolecular ligation efficiency assesses the ligation efficiency of a circularized DNA fragment from Chr.VIII following *EcoR*1 digestion. (**A, E**) Plot of individual values of the HO cutting efficiency of the DLC assay of each strain. (**B, D**) Plot of individual values of the DNA loading control of the DLC assay of each strain. (**C, F**) Plot of individual values of the intramolecular ligation control of the DLC assay of each strain.

**Figure S3. Control experiments for the DLC assay of *htbKR*, as well as for genes that implicated in nucleosome occupancy, examined individually or in double mutant combinations. Related to figure 3.**

The DNA loading control experiment measures the amplification cycle (Cp value) of the control locus, *ARG4*. The efficiency of double-strand break (DSB) induction is determined by comparing the percent amplification from the HO cut site at 2 hours to that at 0 hours. Additionally, the intramolecular ligation efficiency assesses the ligation efficiency of a circularized DNA fragment from Chr.VIII following *EcoR*1 digestion. (**A**) Plot of individual values of the HO cutting efficiency of the DLC assay of each strain. (**B**) Plot of individual values of the DNA loading control of the DLC assay of each strain. (**C**) Plot of individual values of the intramolecular ligation control of the DLC assay of each strain.

**Figure S4. Control experiments for the DLE assays. Related to figure 4**.

The DNA loading control experiment measures the amplification cycle (Cp value) of the control locus, *ARG4*. Additionally, the intramolecular ligation efficiency assesses the ligation efficiency of a circularized DNA fragment from Chr.XII following *Hind*III digestion. (**A**) Plot of individual values of the DNA loading control of the DLE assay of each strain. (**B**) Plot of individual values of the intramolecular ligation control of the DLE assay of each strain.

**Figure S5. The H2Bubi level of each strain, as well as the control experiments for the DLC and DLE assays of *htbKR* and genes implicated in DNA resection, are examined individually or in double mutant combinations. Related to figure 5.**

The DNA loading control experiment measures the amplification cycle (Cp value) of the control locus, *ARG4*. The efficiency of double-strand break (DSB) induction is determined by comparing the percent amplification from the HO cut site at 2 hours to that at 0 hours. Additionally, the intramolecular ligation efficiency assesses the ligation efficiency of a circularized DNA fragment from Chr.VIII following *EcoR*1 digestion. (**A**) Whole-cell extracts were collected by TCA method of each strain and H2Bubi level were subsequent analyzed by Western blot analysis using antibody against histone H2B. A monoubiquitin conjugation to histone H2B results in a band shift to a molecular weight of approximately 25 kDa, marked as H2Bubi. (**B**) DLC signal in WT, *rad9*Δ, *nhp6*ΔΔ and *rad9Δ nhp6*ΔΔ strains. Error bars, SEM (n ≥ 2). *: P< 0.02, **: P= 0.009, ns: not significant. (**C**) Plot of individual values of the HO cutting efficiency of the DLC assay of each strain. (**D**) Plot of individual values of the DNA loading control of the DLC assay of each strain. (**E**) Plot of individual values of the intramolecular ligation control of the DLC assay of each strain. (**F**) Plot of individual values of the DNA loading control of the DLE assay of each strain. (**G**) Plot of individual values of the intramolecular ligation control of the DLE assay of each strain.

**Figure S6. Control experiments for gene conversion assay. Related to figure 6**.

**(A)** In the G2 phase of the cell cycle, only colonies with both double-strand breaks (DSBs) cut was measured. These events resulted in the formation of white colonies (where both strands are repaired by short-tract gene conversion), red colonies (where both strands are repaired by long-tract gene conversion), or sectored colonies (where one strand is repaired by short-tract gene conversion and the other by long-tract gene conversion). (**B**) Distribution of recombinant colony types among the specified strains. Error bars, SEM (n ≥ 2). (**C**) Percentage of occurrences for each strain within the white colonies. Error bars, SEM (n ≥ 2).

**Figure S7. Summary of the DLC and DLE results. Related to figure 7**.

**(A)** Heat map summary of the DLC and DLE values (fold over WT) across all mutants tested in this study. (**B**) Schematic representation summarizing the factors influencing nascent D-loop formation and extension.

## Notes

### Competing Interest Statement

The authors have declared no competing interest.

