## Supplemental Files for "Multifaceted roles of H2B mono-ubiquitylation in D-loop metabolism during homologous recombination repair"

**Figure S1. Related to Figure 1**

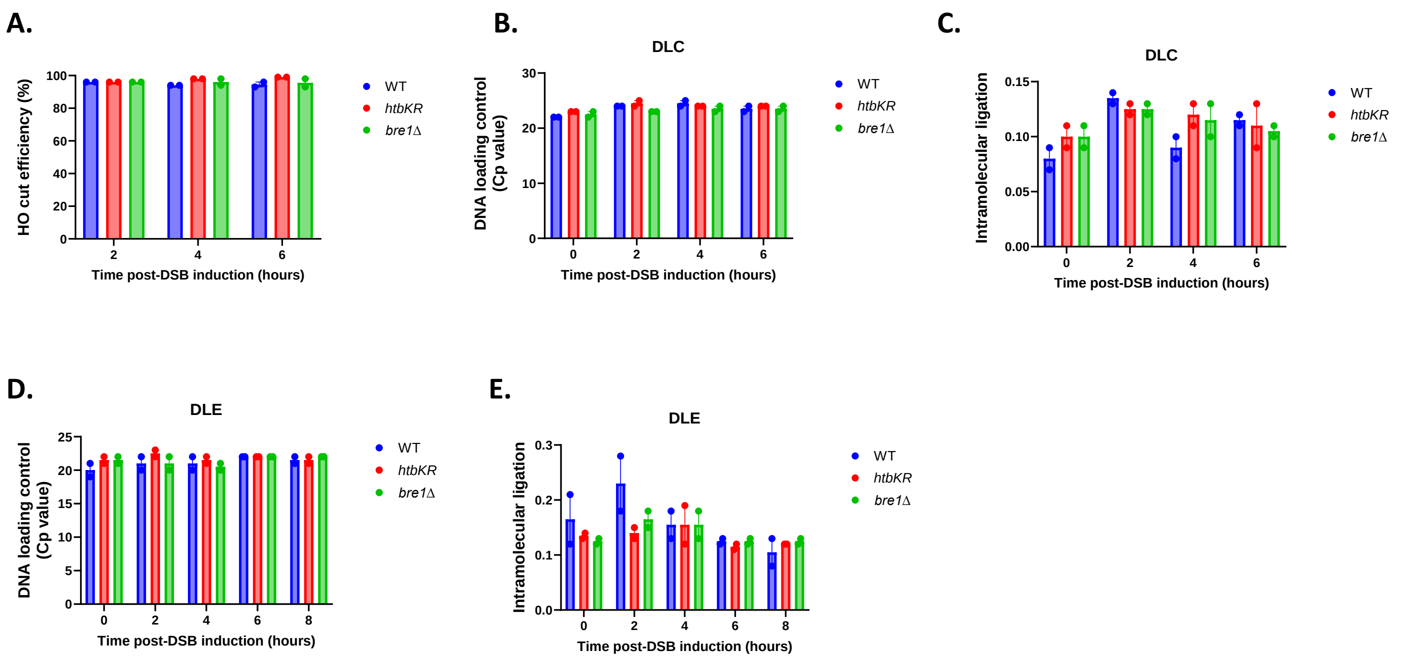

#### Figure S2. Related to Figure 2

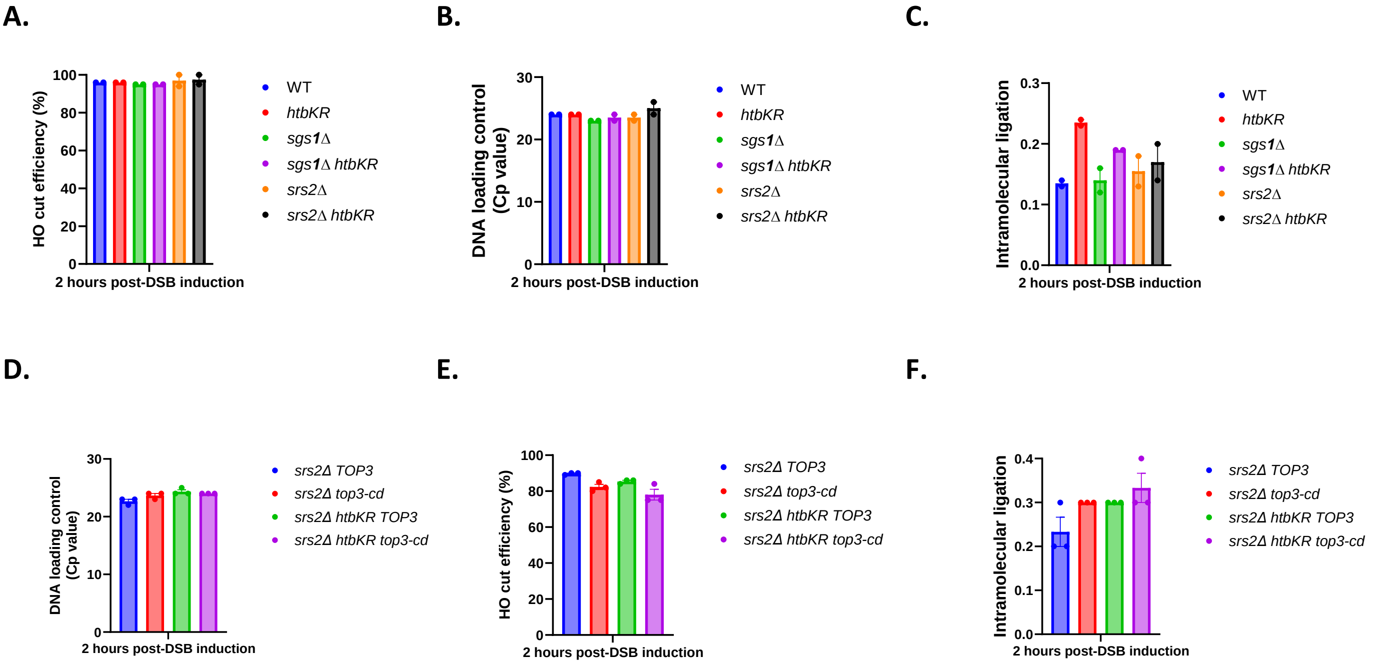

**Figure S3. Related to Figure 3**

**A.**

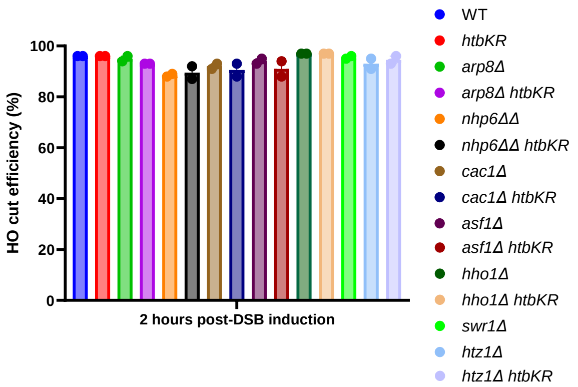


**B.**

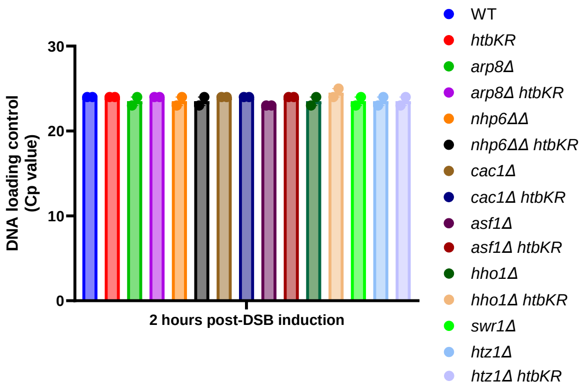


**C.**

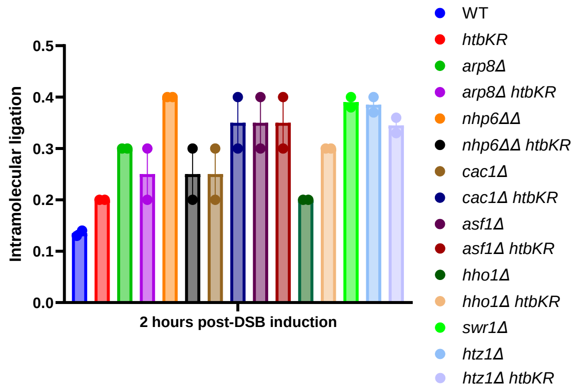

#### Figure S4. Related to Figure 4

**A.**

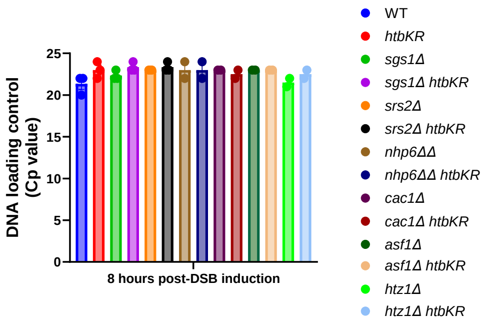


**B.**

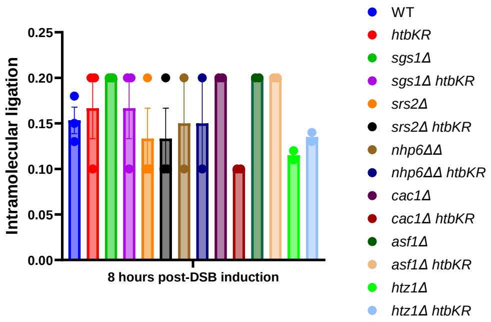

#### Figure S5. Related to Figure 5

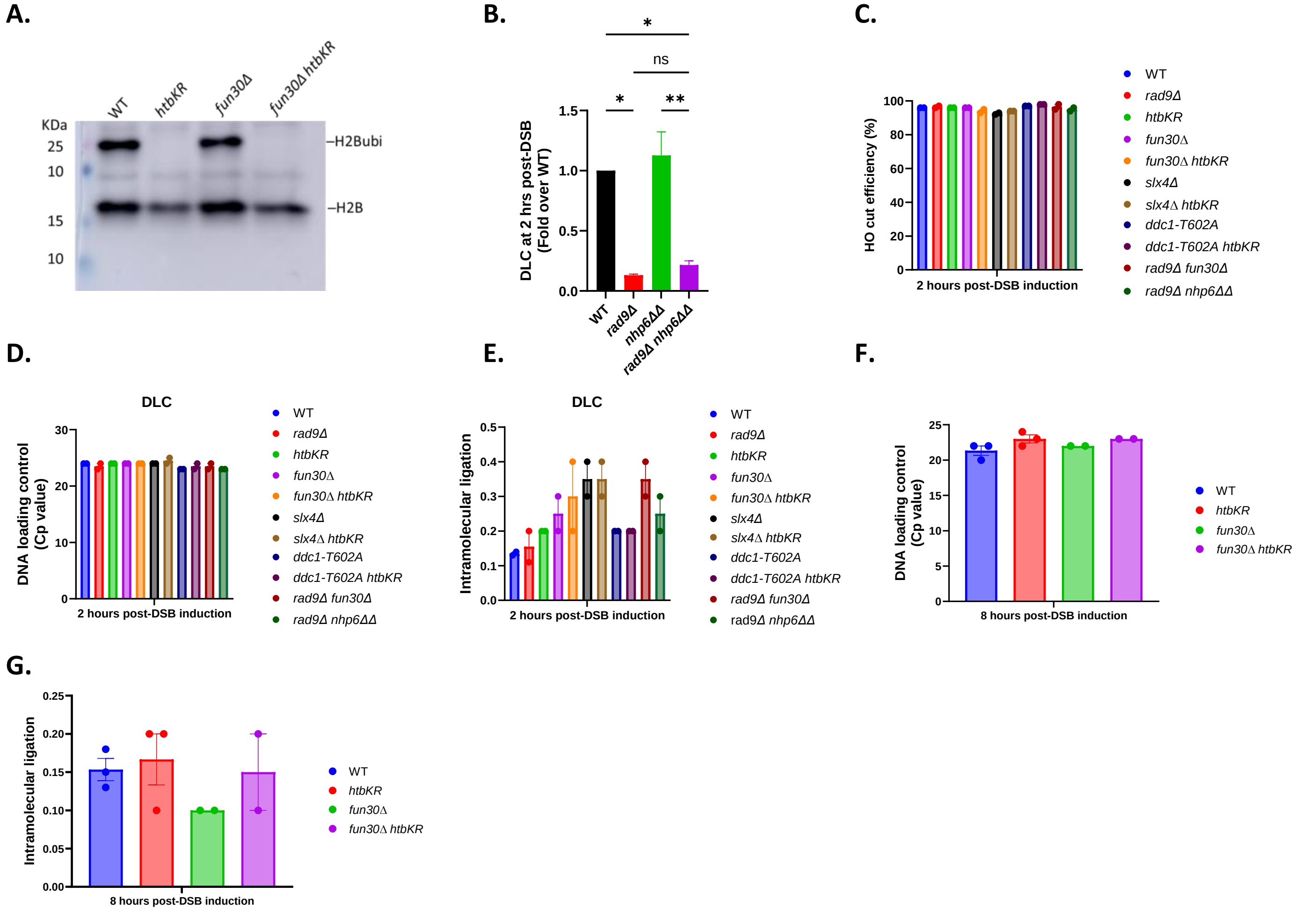

**Figure S6. Related to Figure 6**

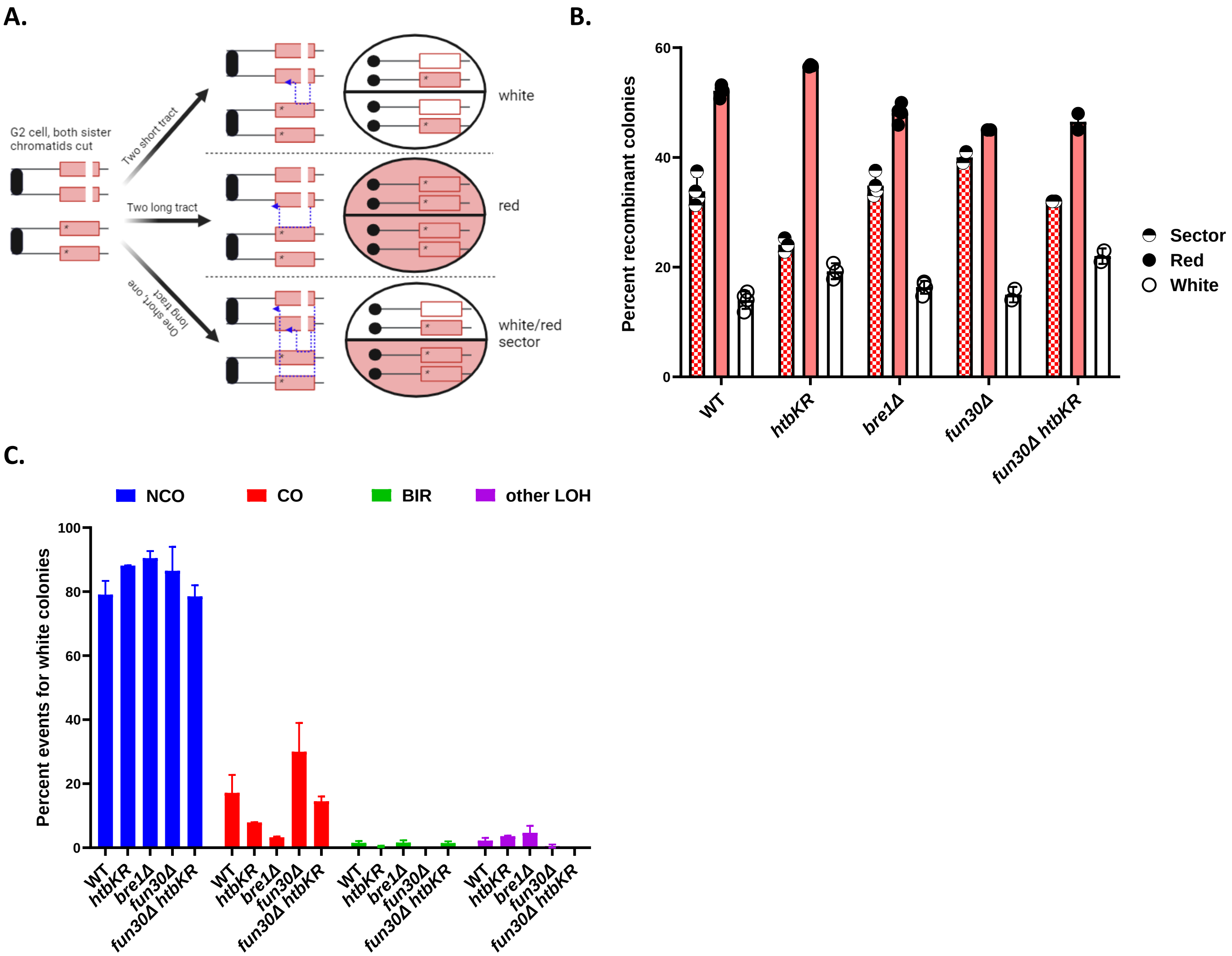

### Figure S7. Related to Figure 7

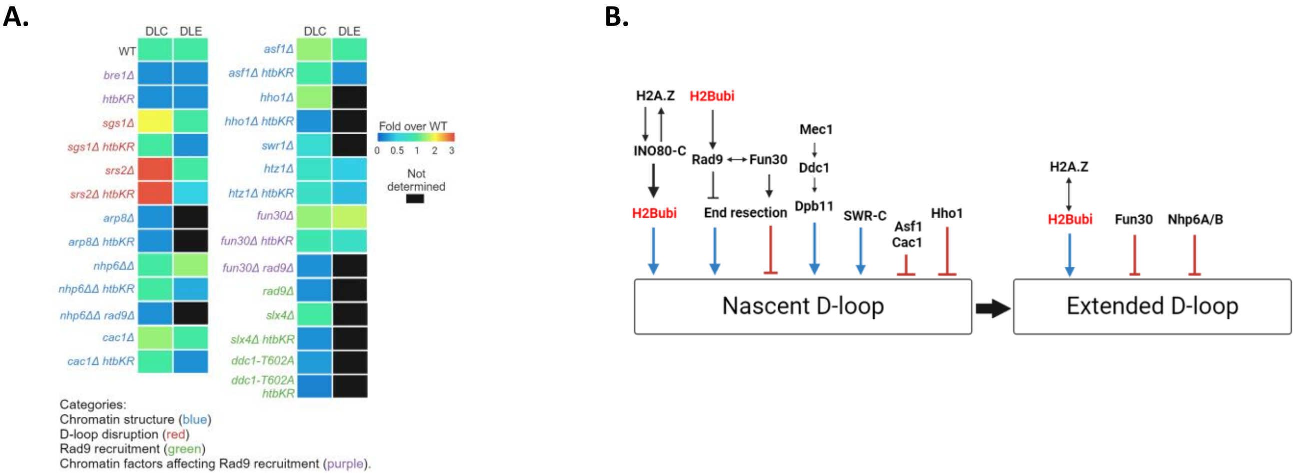

### Table S1. Related to Figure 6

| Summary of DSB-induced recombination events | | | Strain | | | | |
|---|---|---|---|---|---|---|---|
| | | | WT | *bre1Δ* | *htbKR* | *fun30Δ* | *fun30Δ htbKR* |
| White/Red sectored colonies | BIR | $Hph^SNat^R/Hph^RNat^R$ | 8 | 26 | 13 | 4 | 4 |
| | | $Hph^RNat^R/Hph^SNat^R$ | 2 | 4 | 3 | 1 | 3 |
| | | $Hph^SNat^R/Hph^SNat^R$ | 0 | 1 | 0 | 0 | 0 |
| | | **Total no. of BIR** | 10 | 31 | 16 | 5 | 7 |
| | BIR-Like | $Hph^RNat^S/Hph^RNat^R$ | 11 | 19 | 10 | 6 | 3 |
| | | $Hph^RNat^R/Hph^RNat^S$ | 0 | 0 | 2 | 1 | 5 |
| | | $Hph^RNat^S/Hph^RNat^S$ | 0 | 1 | 0 | 0 | 0 |
| | | **Total no. of BIR-like** | 11 | 20 | 12 | 7 | 8 |
| | CO | $Hph^SNat^R/Hph^RNat^S$ | 40 | 22 | 10 | 15 | 27 |
| | | $Hph^RNat^S/Hph^SNat^R$ | 81 | 40 | 34 | 24 | 19 |
| | | **Total no. of CO** | 121 | 62 | 44 | 39 | 46 |
| | **Total no. of NCO** | | 353 | 294 | 187 | 149 | 182 |
| | Chromosome loss | | 0 | 0 | 0 | 0 | 0 |
| | **Colonies analyzed** | | 495 | 407 | 259 | 200 | 243 |
| | Total no.of LOH | | 142 | 113 | 72 | 51 | 61 |
| Red colonies (recombinants) | Sectored LOH | $Hph^SNat^R/Hph^RNat^R$ (BIR) | 21 | 18 | 45 | 2 | 6 |
| | | $Hph^RNat^S/Hph^RNat^R$ (BIR-Like) | 9 | 16 | 12 | 1 | 6 |
| | CO | $Hph^SNat^R/Hph^RNat^S$ | 188 | 63 | 71 | 36 | 42 |
| | Unclassified LOH | $Hph^SNat^R$ | 12 | 30 | 36 | 10 | 26 |
| | | $Hph^RNat^S$ | 1 | 4 | 2 | 4 | 0 |
| | | Total no. of Unclassified LOH | 13 | 34 | 38 | 14 | 26 |
| | Total no. of NCO | | 549 | 416 | 436 | 169 | 273 |
| | Chromosome Loss | | 0 | 0 | 0 | 0 | 0 |
| | **Colonies analyzed** | | 780 | 547 | 602 | 222 | 353 |
| | Total no. of LOH | | 231 | 131 | 166 | 53 | 80 |
| White colonies | Sectored LOH | $Hph^SNat^R/Hph^RNat^R$ (BIR) | 0 | 1 | 1 | 0 | 1 |
| | | $Hph^RNat^S/Hph^RNat^R$ (BIR-Like) | 3 | 2 | 0 | 0 | 2 |
| | CO | $Hph^SNat^R/Hph^RNat^S$ | 19 | 3 | 8 | 5 | 12 |
| | Unclassified LOH | $Hph^SNat^R$ | 0 | 5 | 3 | 1 | 5 |
| | | $Hph^RNat^S$ | 5 | 5 | 4 | 0 | 2 |
| | | Total no. of Unclassified LOH | 5 | 10 | 7 | 1 | 7 |
| | Total no. of NCO | | 187 | 171 | 183 | 70 | 129 |
| | Chromosome Loss | | 0 | 0 | 0 | 0 | 0 |
| | **Colonies analyzed** | | 214 | 187 | 199 | 76 | 151 |
| | Total no. of LOH | | 27 | 16 | 16 | 6 | 22 |

#### Table S2.

| Strain | Genotype | Resource |
|---|---|---|
| WDHY5509 | MATa-inc, ura3::PhiX-A-HOcs, lys2::A, trp1::GAL-HO-hphMX, his3-11,15, can1-100, leu2-3,112, ade2-1, RAD5 | Piazza 2019 Mol Cell |
| WDHY5511 | MATα-inc, ura3::PhiX-A-HOcs, lys2::A, trp1::GAL-HO-hphMX, his3-11,15, can1-100, leu2-3,112, ade2-1, RAD5 | Piazza 2019 Mol Cell |
| WDHY6103 | WDHY5509, bre1::LoxP-kanMX-LoxP | This study |
| WDHY6100 | WDHY5511, htb1-K123R-natMX, htb2-K123R-HIS3 | This study |
| WDHY6224 | MATa-inc, lys2::NatMX4 AVT2::lys-HOcs:: KanMX6 ade3::GAL-H0 URA3::TK C0S9::TRP1-ys2 Ch XI 15 kb donor | Donnianni 2013 PNAS |
| WDHY6225 | WDHY6224, htb1-K123R-natMX, htb2-K123R-HPH | This study |
| WDHY6141 | WDHY5509, sgs1::kanMX | This study |
| WDHY6142 | WDHY5511, htb1-K123R-natMX, htb2-K123R-HIS3, sgs1::KanMX | This study |
| WDHY6143 | WDHY5509, srs2::kanMX | This study |
| WDHY6144 | WDHY5511, htb1-K123R-natMX, htb2-K123R-HIS3, srs2::KanMX | This study |
| WDHY6158 | WDHY5509, srs2::kanMX pYES2 (URA3) | This study |
| WDHY6159 | WDHY5509, srs2::kanMX pYES2-TOP3(URA3) | This study |
| WDHY6160 | WDHY5509, srs2::kanMX pYES2-top3Y356F(URA3) | This study |
| WDHY6161 | WDHY5511, htb1-K123R-natMX, htb2-K123R-HIS3, srs2::kanMX pYES2 (URA3) | This study |
| WDHY6162 | WDHY5511, htb1-K123R-natMX, htb2-K123R-HIS3, srs2::kanMX pYES2-TOP3 (URA3) | This study |
| WDHY6163 | WDHY5511, htb1-K123R-natMX, htb2-K123R-HIS3, srs2::kanMX pYES2-top3Y356F(URA3) | This study |
| WDHY6185 | WDHY5509, arp8::LoxP-kanMX-LoxP | This study |
| WDHY6186 | WDHY5509, htb1-K123R-natMX, htb2-K123R-HIS3, arp8::LoxP-kanMX-LoxP | This study |
| WDHY6180 | WDHY5509, nhp6a::LoxP-kanMX-LoxP, nhp6b::LoxP-kanMX-LoxP | This study |
| WDHY6181 | WDHY5509, htb1-K123R-natMX, htb2-K123R-HIS3, nhp6a::LoxP-kanMX-LoxP, nhp6b::LoxP-kanMX-LoxP | This study |
| WDHY6200 | WDHY5509, asf1::LoxP-kanMX-LoxP | This study |
| WDHY6201 | WDHY5509, htb1-K123R-natMX, htb2-K123R-HIS3, asf1::LoxP-kanMX-LoxP | This study |
| WDHY6198 | WDHY5509, cac1::LoxP-kanMX-LoxP | This study |
| WDHY6199 | WDHY5511, htb1-K123R-natMX, htb2-K123R-HIS3, cac1::LoxP-kanMX-LoxP | This study |
| WDHY6182 | WDHY5509, hho1::LoxP-kanMX-LoxP | This study |
| WDHY6183 | WDHY5511, htb1-K123R-natMX, htb2-K123R-HIS3, hho1::LoxP-kanMX-LoxP | This study |
| WDHY6191 | WDHY5509, swr1::LoxP-kanMX-LoxP | This study |
| WDHY6187 | WDHY5509, fun30::natMX | This study |
| WDHY6188 | WDHY5509, htb1-K123R-natMX, htb2-K123R-HIS3, fun30::natMX | This study |
| WDHY6226 | WDHY5509, slx4::LoxP-kanMX-LoxP | This study |
| WDHY6227 | WDHY5511, htb1-K123R-natMX, htb2-K123R-HIS3, slx4::LoxP-kanMX-LoxP | This study |
| WDHY6106 | WDHY5509, ddc1-T602A | This study |
| WDHY6107 | WDHY5511, htb1-K123R-natMX, htb2-K123R-HIS3, ddc1-T602A | This study |
| WDHY6192 | WDHY5509, htz1::LoxP-kanMX-LoxP | This study |
| WDHY6212 | WDHY5511, htb1-K123R-natMX, htb2-K123R-HIS3, htz1::LoxP-kanMX-LoxP | This study |
| WDHY6207 | WDHY5511, nhp6a::LoxP-kanMX-LoxP, nhp6b::LoxP-kanMX-LoxP rad9::LoxP-kanMX-LoxP | This study |
| WDHY6208 | WDHY5509, rad9::LoxP-kanMX-LoxP fun30::NAT | This study |
| WDHY6020 | WDHY5509, rad9::LoxP-kanMX-LoxP | This study |
| WDHY4357 | MATa-inc, ura3::HOcs (V) lys2::ura3-HOcs-inc (5.6kb), ade3::GALHO leu2-3,112 his3-11,15 trp1-1 ade2-1 | Aylon 2003 MCB |
| WDHY6091 | WDHY4357, htb1-K123R-natMX, htb2-K123R-HPH | This study |
| WDHY6228 | WDHY4357, fun30::Loxp-KanMX-Loxp | This study |
| WDHY6229 | WDHY4357, htb1-K123R-natMX, htb2-K123R-HPH, fun30::Loxp-KanMX-Loxp | This study |
| WDHY6230 | MATα-inc, ade2-IsceIcs, MET22, his3::hphMX4, LSY2, RAD5 | This study |
| WDHY6231 | MATa-inc, ade2-n, met22::KLURA3, his3::NATMX4, lys2::GAL-ISceI, RAD5 | This study |
| WDHY6232 | MATa-inc, ade2-IsceIcs, MET22, his3::hphMX4, LSY2, RAD5, htb1-K123R::LoxP-kanMX-LoxP, htb2-K123R::HIS3 | This study |
| WDHY6233 | MATα-inc, ade2-IsceIcs, MET22, his3::hphMX4, LSY2, RAD5, htb1-K123R::LoxP-kanMX-LoxP, htb2-K123R::HIS3 | This study |
| WDHY6234 | MATa-inc, ade2-I lys2::GAL-ISCEI his3::HPHMX4 bre1::Loxp-KAN-Loxp | This study |
| WDHY6235 | MATα-inc, ade2-n his3::NATMX4 met22::klURA3 bre1::Loxp-KAN-Loxp LYS2 | This study |
| WDHY6236 | MATα-inc, ade2-IsceIcs, MET22, his3::hphMX4, LSY2, RAD5, fun30::LoxP-kanMX-LoxP | This study |
| WDHY6237 | MATa-inc, ade2-n, met22::KLURA3, his3::NATMX4, lys2::GAL-ISceI, RAD5, fun30::LoxP-kanMX-LoxP | This study |
| WDHY6238 | MATa-inc, ade2-IsceIcs, MET22, his3::hphMX4, LSY2, RAD5, htb1-K123R::LoxP-kanMX-LoxP, htb2-K123R::HIS3, fun30::LoxP-kanMX-LoxP | This study |
| WDHY6239 | MATα-inc, ade2-n, met22::KLURA3, his3::NATMX4, lys2::GAL-ISceI, RAD5, htb1-K123R::LoxP-kanMX-LoxP, htb2-K123R::HIS3, fun30::LoxP-kanMX-LoxP | This study |
